# Stereo-EEG reveals rich cortical dynamics in humans coping with difficult action discrimination

**DOI:** 10.1101/2020.01.23.916569

**Authors:** A. Platonov, V. Pelliccia, I. Sartori, G. LoRusso, P. Avanzini, G.A. Orban

## Abstract

Visual perception of others’ actions is important for social interactions, and the ability to do so, even when one gets only brief glimpses of others’ behavior, may be crucial for survival. At present it is unknown how the human brain solves this problem. Imaging studies have promoted the idea that the multiple demand (MD) system, a domain general system of the human brain, operates in difficult cognitive and perceptual tasks, but not in tasks in which sensory information is reduced. Functional imaging, based on slow hemodynamic responses, may miss or standardize neural events with very diverse time courses. Here we exploited the temporal resolution of stereo-EEG to study directly cortical activity when human subjects must judge the actions of others, but only get brief glimpses of others’ activity, because the videos were truncated. Multiple cortical regions increased their activity in the difficult action discrimination, relative to the easy task when the complete video was visible. The majority of these regions belonged to the MD system, being located in parietal or prefrontal cortex. The variety of time courses, lasting from a few 100ms to several seconds, allowed us to disentangle control from effector regions, the latter processing observed actions. This distinction was further supported by relationships with behavior. A key operation within the control clusters was the prediction of erroneous responses, which was initiated in the PPC soon after the end of the truncated video. The time courses further suggested that MD regions not only exert control, but also perform various evaluations of the effort, important for efficient and intelligent behavior. We observed also increases outside the MD system, in temporo-parietal cortex, which may provide contextual information about variables related to the observed action, such as the actor, the object or the scene. Furthermore, to cope with the brief sensory input, the MD system called upon warning regions reacting to the static picture of the actor presented just before the video. We conclude that discrimination of brief observed actions indeed involves the MD system, which is thus is more general than assumed so far. WE also show that the MD system is more complex than assumed, as it includes evaluation of control, and more flexible, as it interacts with other systems than simply the effector circuit of the task.

## Introduction

Social interactions belong to the core of human behavior, underscoring the importance of perceptual mechanisms processing visual signals arising from others actions (Platonov et al 2016, 2017). Such processes are particularly challenged when subjects get only brief glimpses of others’ behavior, as when fighting an enemy hiding in the environment. The present study investigates how the human brain copes with such a difficult task, critical for human survival. When we get only brief glimpses of the other’s actions, the difficulty arises from a reduction in visual information (Norman & Bobrow 1975), which can be severe, as duration thresholds of action observation are quite short (Tucciarelli et al 2015, Platonov and Orban 2016). At present it is unknown how the brain solves a task when difficulty arises from sensory limitations. The multiple demand (MD, Duncan 2010), or cognitive control system, is a domain general system in parietal and prefrontal cortex, active when subjects face difficult cognitive tasks (Duncan & Owen 2000, Fedorenko et al 2013). Initially, the MD system was considered monolithic and operating in all difficult cognitive tasks (Crittenden & Duncan 2014), but more recently (Crittenden et al 2016) it was shown to operate in perceptual as well as cognitive tasks, and to include lateral prefrontal and cingulo-insular subnetworks, reminiscent of an earlier distinction (Dosenbach et al 2008). The MD system interacts with the networks underlying the various cognitive or perceptual tasks, and its activity reflects the effort (Shenhav et al 2017) devoted by the subjects to appropriately perform those tasks. However, Han and Marois (2013) and Wen et al (2018) have demonstrated that the MD system is not involved in perceptual tasks in which the difficulty resulted from reduced sensory information. This negative result may reflect the low-level of the features used in the discrimination task (direction of moving dots) of Wen et al (2018), but earlier studies of difficult speed discrimination (Sunaert et al 2000) did find prefrontal activations. Furthermore, Han and Marois (2013) used letters as targets in their rapid serial visual presentation task. Alternatively, the negative results of Han and Marois (2013) and Wen et al (2018) may be due to technical limitations, imposed by the low temporal resolution of functional MRI (Ghuman and Martin 2019). Stereo EEG (sEEG) has millisecond temporal resolution, spatial resolution of a few mm (Parvizi and Kastner 2018), and by integrating data from multiple patients (Avanzini et al 2016) adequate coverage of the hemispheres. Therefore, the first aim of the present study was to investigate using sEEG, whether or not the MD system is involved in difficult action discrimination, by comparing cortical activity during action discrimination based on truncated and full videos.

The use of sEEG with its excellent temporal resolution to study difficult action discrimination allowed us to address a second question. Indeed, to understand how the brain copes with difficult action discrimination, we need to document the involvement of the MD system, but also, if it is involved, its interactions with regions processing observed actions. Functional imaging studies have defined the MD system spatially, as consisting of the lateral and orbital parts of the middle frontal gyrus, the opercular part of the inferior frontal gyrus, the anterior insula, the precentral gyrus, supplementary motor cortex, and anterior cingulate in the frontal cortex, as well as inferior and superior parietal cortex (Fedorenko et al 2013). These latter parietal parts overlap with the action observation network (AON, Cross et al 2009), which includes regions at the occipito-temporal, parietal, and premotor levels (Caspers et al 2010, Jastorff et al 2010). Hence to distinguish amongst the regions active in a difficult action discrimination task, those belonging to the MD system or to the AON is difficult using fMRI, because of its low temporal resolution. Thus the second aim of the present study was to investigate, using sEEG, the interactions between the MD system and the action observation regions. The choice of sEEG had the additional benefit of revealing the dynamics of the fronto-parietal regions of the MD system involved in difficult action discrimination, which constituted a third aim of the study. So far, only the dynamics of a few prefrontal MD regions involved in error monitoring (Tang et al 2016), or the encoding of reward values (Saez et al 2018) have been investigated using sEEG.

Our results confirmed the involvement of the MD system and the complexity of its dynamics, including unexpected late responses. Furthermore they showed the recruitment of auxiliary cortical circuits, in addition to the MD system and action observation regions.

## Results

### Behavioral results

Twenty eight patients (15 male, 13 female, age 18-49, mean 32 years) undergoing functional exploration with stereo-EEG recording in view of surgical interventions to cure focal drug resistant epilepsy, gave their informed consent to participate in the present study. The demographic and clinical data of the patients, 24 of which participated in a previous study (Platonov et al 2019), are listed in table S1, and their neuropsychological results in table S2. During testing, the experimenter verified that patients fixated well. This was confirmed by eye position recordings: the grand mean of standard deviations of horizontal and vertical eye position in 18 patients were close to 1deg in all four conditions (Table S3). In particular neither the horizontal (two tailed t test, t= 0.66, df=17, p>0.52) nor the vertical (t = 2.09, df=17, p>0.05) eye position differed significantly between long and short action trials.

Stimuli and tasks were exactly the same as in Platonov et al (2018, see methods). Patients viewing either complete or truncated videos showing an actor performing a manipulative action, discriminated either the observed action (grasp vs push), or the actors’ gender (male vs female). Trials were blocked according to task and duration of videos. Each trial (fig 1) included a 1s baseline period, an interval in which the first frame of video was shown for a variable interval (283 to 883ms), followed by the video presentation (1167ms) and a 2s period during which subjects were allowed to signal their response. Subjects were instructed to respond when ready, and the video ended when the subjects responded. The static frame preceding video onset was identical for the two actions, and thus provided gender but no action information. In the short trials, the video was shown only briefly and replaced during the remaining time by dynamic noise. Hence in these trials, the dynamic period (short video plus dynamic noise) replaced the video period of long trials. Before the recordings, the duration thresholds (84% correct) of action discrimination were determined individually, and the truncated videos in the short trials lasted either the exact duration threshold (228ms on average), 100ms less or 100ms more.

**Fig 1:**
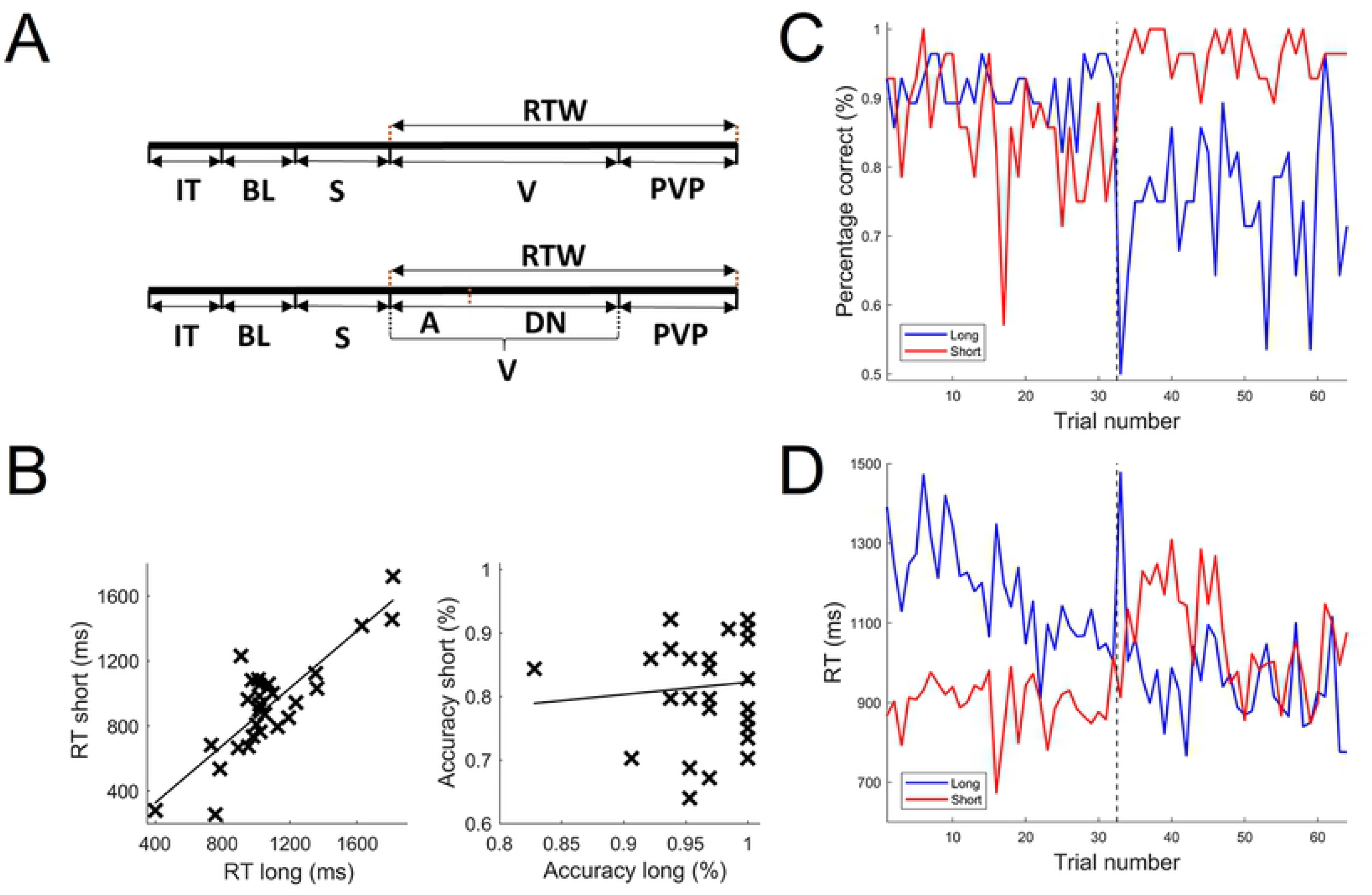
behavioral paradigm and performance: A two types of trials: long (top) and short (bottom) trials B: RT and accuracy of observed action discrimination in short trials as a function of these variables in the long trials; C&D: single trial performance of the population, accuracy (C) and RT (D), in the two action discrimination blocks: the first (blue) starting with 32 long trials followed by 32 short trials, and the second (red) with short and long trial sub-blocks in reverse order. Abbreviations: IT: inter-trial, BL: baseline, S: static, V: video, A: action, DN: dynamic noise, RTW: response time window, PVP: post video period.

In short action trials, accuracy (mean 82%, SD 9%, range 63-95%) was significantly (paired t test, t=8.9, df 27 p<0.001) reduced compared to long action trials (mean 97%, SD 4%, range 83-100). The reaction time (RT), relative to video onset, in short action trails (927ms, SD 297, range 256-1712ms) were also significantly (paired t test t=4.7, df 27, p<0.001) shorter than in long action trials (mean 1100ms, SD 274, range 732-1821ms). The RT for short (RT_s_) and long (RT_l_) trials correlated significantly (r=0.78, p<10-5, eq RT_s_=1.4+0.84RT_l_), indicating a strong patient factor contributing to the RTs. RT and accuracy in short trials also correlated (r=0.43, p<0.05), indicating that as patients responded later their performance improved. One patient with very short RT (256ms) and low accuracy (63%) in short action trials was removed from the analysis of the stereo-EEG data. While the truncation of the videos had a strong effect on action discrimination, it barely affected gender discrimination, which used predominantly the static presentation preceding the video (Platonov et al 2019). RTs, relative to static onset, averaged 871ms (SD 372ms) for long compared to 816ms (SD 355ms) for short trials (paired t test t= 2.7, df 27, p<0.02), and accuracy equaled 97% (SD 4%) and 97% (SD 5%) for long and short gender trials. Thus accuracy was unchanged and RTs shortened modestly and nonspecifically for short gender trials. Hence, the behavioral data indicate that truncation of the video made action discrimination, but not gender discrimination, more difficult.

To investigate how behavior evolved during an action block, and in particular the effects of switching from long to short trials or the reverse, we averaged RT and accuracy across the 27 patients for each of the 64 trials in a block. In the first block (blue curves in fig 1), accuracy was consistently high (close to 90% correct) in the first 32 (long) trials, dropped to 50% at the first short trial, but recovered rapidly, hovering around 75% correct in the remaining short trials. In this block RTs started at large values (around 1300ms) and then steadily decreased until the first short trial, in which they increased to nearly 1500ms, resuming their decrease afterwards, to reach values around 950ms in the last 20 trials. In the second block (red curves in fig 1) changes were more limited: accuracy oscillated around 85% correct during the short trials, and jumped to 95% during the long trials. RTs oscillated around 900ms during the short trials, increased transiently to 1100ms during the long trials, but returned to values around 950ms in the last 20 long trials. Thus patients adapted very rapidly, within two trials, to changes in video duration, and modestly improved in the second compared to the first action block.

In conclusion, the behavioral results show that the patients performed the tasks well and that short action discrimination trials, but not short gender trials, were more difficult than their long counterparts. Turning to the sEEG recordings to characterize cortical activity increases in difficult action discrimination, yielded findings expected for effector and control regions (Fedorenko et al 2013, labeled 1-2 in fig 2), but also three unexpected findings (3-5 in fig 2). Analyzing the wide variety of time courses of the increases, revealed increases during the entire dynamic period but also restricted to the initial part of the dynamic period (fig 2), as expected for control and effector regions respectively. The latter are involved in action discrimination, and should more specifically provide sensory evidence about observed actions (fig 2A), and thus include regions of the action observation network, active even in passive viewing (see supplementary text). However, unexpectedly, a third group of leads increased their activity only at the end of the dynamic period, indicating additional processing in short action trials (4 in fig 2B). Next, clustering the 3 groups of enhanced leads to compute their overlap with the MD system, revealed a third of the leads enhanced over the entire dynamic period to be located outside the MD system, again documenting unexpected additional processing in short action trials (3 in fig 2B). Finally, extending the analysis to the static period preceding the video-onset, we found leads with increased activity during this static period, indicating again additional processing in short action trials (5 in fig 2B). Having documented three types of dynamic period increases as well as static period increases, we investigated the detailed time courses of activity increases in the individual clusters of enhanced leads, to document the properties predicted for control and effector regions, and to provide clues as to the function of unexpected enhanced regions.

**Fig 2:**
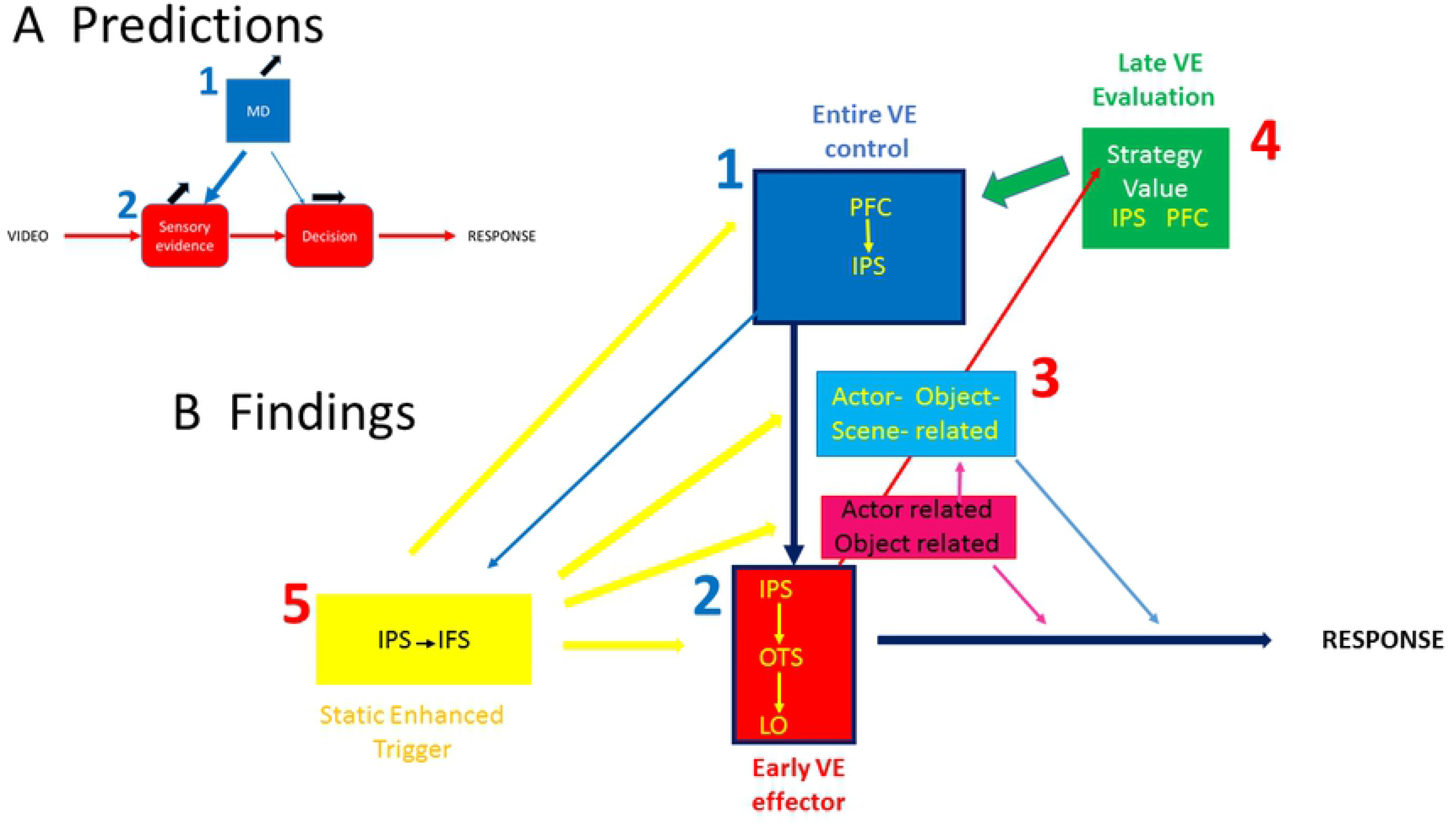
Comparison of predicted and observed increases in difficult action discrimination. A predicted increases in MD and action discrimination circuit, including sensory evidence collecting and accumulating stages; B: schematic diagram of the five (numbers) types of increased regions: the two parts of the MD system (blue and green plain colors), the effector regions (red), and two associated systems: the static trigger (yellow) and the regions providing actor-related and other additional information (light colors). The dark blue outlines indicate the two parts that were expected from previous studies and shown in A. Black arrows in A indicate changes with difficulty; arrows in B possible interactions between regions.

### Clusters of Video enhanced (VE) leads

#### Video-enhanced leads

Stereo-EEG recordings were generally made in a single hemisphere (16 right, 9 left), with 3 bilateral implantations. Thus the right hemisphere was sampled more than the left (fig S1): 2249 of the 3406 leads located in grey matter were attributed to the right hemisphere (19 hemispheres, 118/h), and 1157 to the left (12 hemispheres, 96/h). About half of the recorded cortical leads (1669, 49%) were involved in the action discrimination task, as demonstrated by a 2 (video duration: long and short) x 3 (period: baseline, static, video/dynamic) ANOVA of averaged z-scored gamma activity yielding an FDR-corrected significant main effect of period or a significant interaction (fig S2). The proportions being very similar in the right (1104 leads, 49%) and left (565 leads, 49%) hemispheres, (also for the leads with enhanced activity in short action trials, see below), and the MD system being bilateral (Fedorenko et al 2013), we will concentrate on the right hemisphere, which had better coverage (fig S1).

The large diversity of the time courses of the increase in the dynamic period of the short compared to long trials, required an analysis with flexible time windows. We used three criteria (see methods) to define video-enhanced (VE) leads, two handling the diversity of timings of the increased activity in the short trials, and the third criterion preventing OFF responses to the end of the video to be considered increases in the short trials (OFF-responses were far more common than anticipated, occurring also in high-level visual regions, and will be the topic of a later report). This procedure yielded 432 VE-leads in the right hemisphere and 193 in the left, corresponding the 19% and 17% of the tested leads respectively. The next step was to distinguish (see methods) amongst the VE leads those with an increase during the truncated video itself (early-VE), those showing an increase later in the dynamic period, during the noise presentation (late-VE), and those showing an increase over most of the dynamic period (Entire-VE). About 10% of the VE leads were classified as Early-VE leads, a third as Late-VE and the majority as Entire-VE leads (Table 1).

**Table 1:** distribution of video enhanced leads.

| type | right | Left | Total | % of video enhanced |
| --- | --- | --- | --- | --- |
| Early | 47 | 17 | 64 | 10 |
| Entire | 265 | 97 | 362 | 58 |
| Late | 120 | 79 | 199 | 32 |
| Total | 432 (19% tested) | 193 (17% tested) | 625 (18% tested) | 100 |

The average time courses of these three categories of VE leads confirmed their differences (fig 3): Early-VE leads increased their activity during the end of the static period and the first 500ms of the dynamic period of the short action trials, Entire-VE leads during the end of the static period and the whole of the dynamic period, and Late-VE leads only after 350ms into the dynamic period. While the first two of these time courses correspond to what was expected for the initial steps of the action discrimination circuit, and the multiple demand control system respectively, the late VE leads were the first of our *unpredicted findings* (labelled 4 in fig 2). The time courses also showed that the increases in short trials were specific to action discrimination (fig 3A): long and short trials did not differ in the gender task, with the Early-, Entire-, and Late-VE leads showing a strong, weak and no response to the static presentation, respectively. The lack of difference between short and long gender trials, confirms that the increases in short relative to long action trials reflect difficulty. The proportions of the 3 VE categories were relatively similar in the two hemispheres, the proportion of Late-VE leads being slightly larger, and that of the two other categories slightly less in the left hemisphere (χ^2^ =10.7, p<0.05, Table 1). Also the time courses of the increased activity in the short action trials were similar in the two hemispheres for the 3 categories (fig S3), although the activation of the Early-VE leads, both in short and long action trials, was weaker in the left than the right hemisphere, due to a lesser sampling of occipito-parietal regions in the left hemisphere..

**Fig 3:**
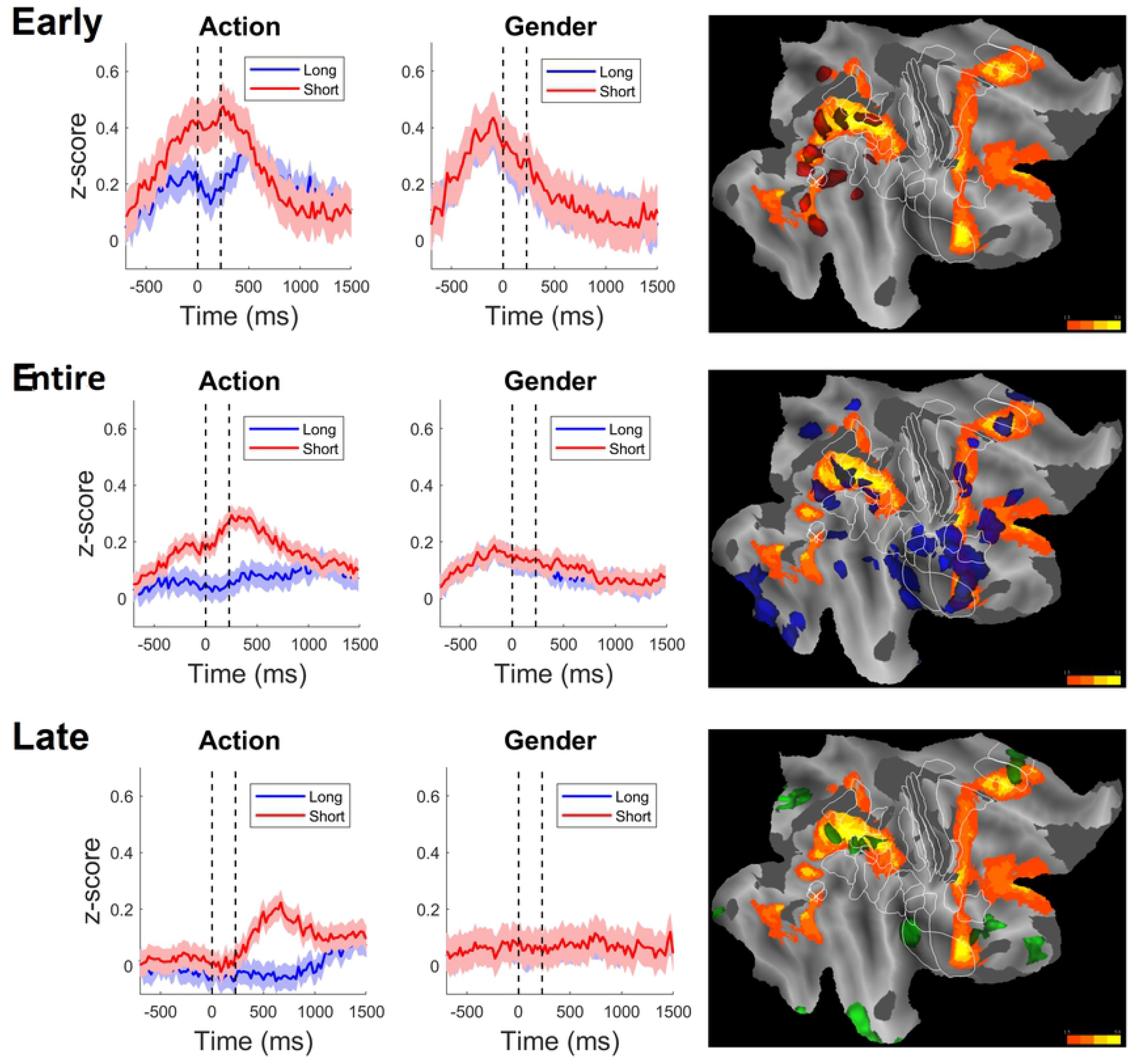
Three types of Video Enhanced (VE) clusters: the time courses of z-scored gamma power in the four conditions (left), action and gender long (blue) and short (red), and the localization on flatmaps of the right hemisphere (right) for early video enhanced (red, top), entire video enhanced (blue, middle), and late video enhanced (green, bottom). Orange regions MD system as defined by Fedorenko et al, grey hatching underexplored regions (see fig S1). Vertical dotted lines indicate average video duration in short trials. Hatching indicates SE. Color code: 15% to 50% VE-leads. Note the task specificity of the activity increases with difficulty. For time courses in the four conditions for left and right hemisphere separately see fig S3.

#### Definition of VE clusters

The VE leads occurred throughout cortex, but the Early-VE leads occurred primarily in occipito-parietal cortex, while the Entire-VE leads, and even more so the Late-VE leads, were generally absent from occipital cortex. Furthermore, while leads were scattered across the cortical surface, conspicuous accumulation appeared in certain cortical regions, eg Entire-VE leads at the level of the anterior insula,, and inferior frontal gyrus or the Late-VE leads on the medial side of the hemisphere at the level of the anterior Middle cingulate cortex (aMCC, Caruana et al 2017), To clarify the concentration of VE leads of a given category, and to investigate their location with the respect to the MD system, we clustered the VE leads using small circular masks (see methods). This procedure yielded 9 small Early-VE clusters, all located in occipital, parietal or posterior temporal cortex (fig S4). To enhance the representability, small neighboring clusters were grouped together: the two small clusters in anterior and posterior occipito-temporal sulcus (OTS) into a single OTS cluster, the 3 small clusters in posterior, middle and anterior intraparietal sulcus (IPS) into a single IPS cluster, and the small posterior Middle Temporal Gyrus (pMTG) and posterior Superior temporal sulcus (pSTS) clusters into a single posterior temporal cortex (pTC) cluster. Together with the Lateral occipital (LO) cluster and the parieto-occipital sulcus (POS) cluster, this yielded 5 Early-VE clusters, each with leads from 2 to 5 patients (fig S4, Table 2). These clusters included 29 of the 47 Early-VE leads (62%) in the right hemisphere. A similar procedure yielded 22 clusters of Entire-VE leads, with 3 IPS clusters, but those included many leads from several patients and were kept separate. However, three hippocampal clusters were grouped, two pMTG clusters, and one cluster at the border between the frontal operculum (FO) and anterior insula with the anterior insula (aIns). This regrouping yielded 18 Entire-VE clusters (fig S4): 4 in parietal cortex, 4 in temporal cortex and 10 in prefrontal cortex, which together included 188 /263 of Entire-VE leads (71%). All 18 clusters, except dorsal premotor cortex (PMd), included leads from at least 3 patients (Table 2). Finally the clustering yielded 8 Late-VE clusters, one in anterior temporal lobe, one in posterior insula, one in the parieto-occipital sulcus (POS), extending into the calcarine sulcus, and the remaining 5 in prefrontal and parietal cortex (fig S4). No further regrouping was made, and except for ATL, the Late-VE clusters included leads from at least 3 patients (Table 2). Together the 8 clusters included 55 out of 120 Late-VE leads (46%).

**Table 2:** characteristics of the VE clusters.

| Class | RegionName | # of patients | # of leads | AvgMD | N of EZ leads |
| --- | --- | --- | --- | --- | --- |
| Early | LO | 3 | 7 | 0.76 | 1 |
|  | OTS | 2 | 4 | 2.13 | 0 |
|  | pTC | 5 | 6 | -0.55 | 2 |
|  | IPS | 5 | 10 | 3.38 | 0 |
|  | POS | 2 | 2 | 0.65 | 0 |
| Entire | PHIP | 7 | 29 | -0.11 | 8 |
|  | POS | 3 | 6 | -0.08 | 0 |
|  | preC | 3 | 4 | 0.47 | 0 |
|  | pIPS | 7 | 15 | 1.59 | 0 |
|  | mIPS | 4 | 6 | 3.04 | 0 |
|  | aIPS | 5 | 7 | 2.23 | 0 |
|  | pMTG | 4 | 8 | -0.32 | 1 |
|  | pSTS | 3 | 6 | -0.86 | 2 |
|  | TO | 8 | 15 | -0.89 | 0 |
|  | OP1-4 | 5 | 12 | -1.85 | 0 |
|  | BA44 | 7 | 14 | 0.48 | 1 |
|  | FO | 8 | 20 | 1.18 | 0 |
|  | aIns | 5 | 11 | 2.89 | 2 |
|  | IFS | 6 | 13 | 2.05 | 1 |
|  | mPreCG | 4 | 8 | 2.27 | 0 |
|  | PMd | 2 | 3 | 1.73 | 0 |
|  | aMCC | 4 | 8 | 2.61 | 2 |
|  | aSFS | 3 | 3 | 1.57 | 1 |
| Late | ATL | 2 | 8 | 0.00 | 5 |
|  | aFO | 3 | 7 | 0.74 | 4 |
|  | FO1 | 3 | 7 | 0.12 | 0 |
|  | pInsula | 4 | 6 | -0.91 | 0 |
|  | aMCC | 3 | 5 | 1.86 | 1 |
|  | pIPS | 3 | 3 | 3.52 | 0 |
|  | aIPS | 5 | 8 | 2.60 | 0 |
|  | POS | 5 | 11 | 0.78 | 0 |

For distribution of MD values, see fig S5.

#### Overlap with the MD system

The overlap of these VE clusters with the MD system was computed by the averaging t scores in the MD mask of the leads in each cluster (fig 3). Clusters with an average t value exceeding 1.5 (criterion used in Fedorenko et al 2013) were considered belonging to the core of the MD system. Clusters of which 15% of the leads (min 1/7=16%) had a value exceeding 1.5, were considered as being on the edge of MD system: all these clusters had a strictly positive average t value. The other clusters were considered outside the MD system (with one exception their average t-score was zero or less). Thus all clusters considered belonging to the MD system (core and edge combined) had positive average t scores, the lowest positive value being 0.12 in the orbito-frontal cortex (OFC, FO1,Table 2). Three of the five Early-VE clusters belonged to the MD system, with the IPS and OTS clusters belonging to the core (Table 2, fig S5) and the LO cluster to the edge. Twelve of the 18 Entire-VE clusters, all located in parietal or prefrontal cortex, belonged to the MD system, with 9 clusters belonging to the core: posterior IPS (pIPS), middle IPS (mIPS), anterior IPS (aIPS), anterior Insula (aIns), inferior frontal sulcus (IFS), middle precentral gyrus (mPreCG), PMd, anterior Superior frontal sulcus (aSFS) and aMCC, and 3 to the edge: BA44, FO and Prec. Finally, 6 of the 8 Late-VE clusters belonged to the MD system, with the pIPS, aIPS and aMCC belonging to the core, and FO1, aFO and POS to the edge (fig S5, Table 2). Although the proportion of clusters belonging to the core of the MD was slightly larger amongst the Entire-VE clusters than the two other types, the association was not significant (χ2= 1.54, p>0.8). Control analyses (see supplementary information) indicated that by concentrating on the clustered VE leads we described the vast majority (103/126, 82%) of the leads belonging to the MD system.

#### Overlap with the EZ

Finally, we investigated the overlap of the VE clusters with the Epileptic zone (EZ). Only 29 leads of the VE clusters were localized in the EZ: 3 leads in early-VE clusters, 18 in Entire-VE clusters, and 8 in the Late-VE clusters. Only two Late-VE clusters were removed because of excessive overlap with the EZ, as was one late-VE cluster with a spurious increase reflecting the shorter RT (see supplementary text). The 28 remaining VE clusters in the right hemisphere were included in the analysis: 5 Early-VE clusters, 18 Entire-VE clusters, and 5 Late-VE clusters. Of these 28 clusters, 14 belonged to the core of the MD system, 6 to the edge of the MD system, and 8 were outside the MD system. Those belonging to the core of the MD system were, with one exception, all located in parietal (around the IPS) and prefrontal cortex. All clusters outside the MD system were located in temporal cortex or the edges of parietal cortex.

### Clusters of Static Enhanced (SE) leads

The Early and Entire-VE leads enhanced their activity almost immediately after video onset, some even slightly before the onset of the brief video, raising the question of how the brain knows it has to enhance these activities. One possibility are tonic signals during the short action subblocks (Han & Marois 2013). We looked for differences in the baseline activity specifically before the short action trials by calculating the interaction between task and video duration in a 2×2 ANOVA and failed to find any significant interaction surviving FDR-correction. This is not surprising as no explicit instructions were given concerning short trials and patients adapted rapidly to the change in duration of the videos (fig 1). In the absence of tonic signals during the short action subblocks, the Early- and Entire-VE clusters need a trigger signal indicating in each trial that the brief video is about to start, in order to increase their activity in time. The static stimulus period preceding video-onset contained no information about the upcoming action, but provided a robust warning signal by its systematic association with the video. We therefore looked for leads increasing their activity in the static period of the short action trials. Forty nine static-enhanced (SE) leads (39 in right hemisphere, 10 in left) showed a significant increase during the static period, representing yet another *unpredicted finding* (labeled 5 in fig 2). The SE leads represented only 1.5% of the tested leads, with a slight asymmetry favoring the right hemisphere (χ^2^ = 3.49, p<0.062). Their activity (fig 4) in short action trials was enhanced in the static and less so in the beginning of the dynamic period, but for a shorter time than in the Early VE leads. Again the enhancement was completely specific, the leads responding equally in the static period of short and long gender trails. We tested for leads with both static and video enhancement, and found that 2 SE leads were also Early-VE, 16 Entire-VE and 6 Late-VE. Thus the proportion of SE leads was larger amongst VE leads than the overall population of tested leads, reaching 5% amongst the Entire-VE leads. The association of SE leads with Entire-VE leads was significant (χ^2^= 4.3, p<0.05), but not that with Late VE leads (χ^2^ =2.5, ns), the numbers of Early VE leads being too small to test statistically.

**Fig 4:**
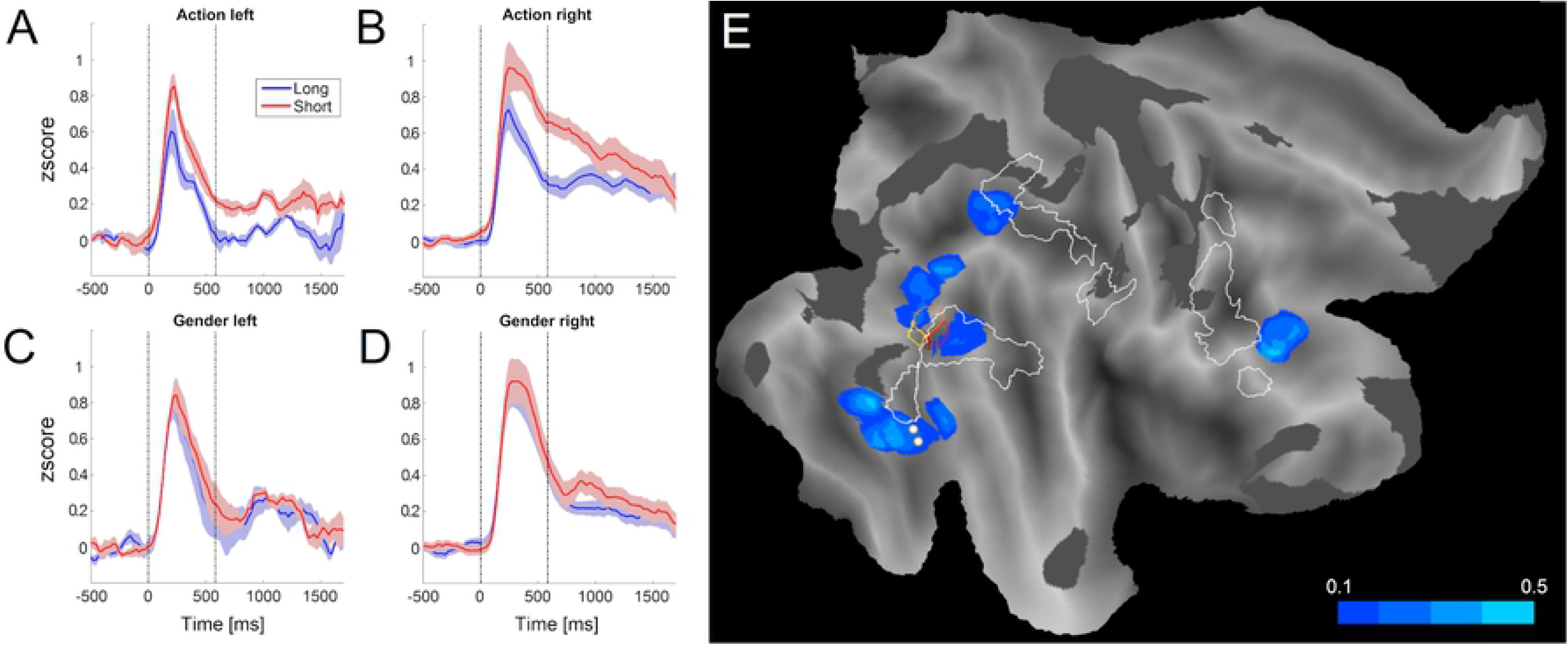
Static enhanced (SE) leads: A-D: The time course of activity in the four conditions action (A, B) and gender (C, D) long (blue) and short (red) for left (A,C) and right (B,D) SE leads. Vertical dotted lines indicate average static period. E:flatmap of right hemisphere showing the five clusters of static enhanced leads (blue) Colored outlines MT retinotopic cluster from Abdollahi et al (2014), white outlines: action observation network from Caspers et al 2010; white dots Local maxima of FBA (Peelen et al 2005); grey hatching underexplored regions (fig S1). Color code: 10 to 50 % SE-Leads. Note the task specificity of the increases with difficulty. Note also that the MT cluster separates the LO and MTG components of EBA.

Clustering of the SE leads (see methods) yielded 5 clusters (fig 4) which included 22 (56%) of the 40 SE leads in the right hemisphere. None of these clusters had leads located in the EZ (Table 3). Two clusters (in LO and in MTG.) were attributed to a single region, the extrastriate body area (EBA), being located on either side of the middle Temporal (MT) retinotopic cluster, which is the hallmark of EBA (Weiner & Grill-Spector 2011), A fusiform gyrus (FG) cluster corresponded to the fusiform body area (FBA, Orlov et al 2010, Peelen & Downing 2005) but extended into the neighboring fusiform face area (FFA, Kanwisher et al 1997). Finally, one cluster was located in the IPS, more dorsal to a body region described by Orlov et al 2010), and another in the IFS, in the prefrontal face area described by Tsao et al (2008). Three SE clusters belonged to the core of the MD system (IPS, IFS, and LO), one to the edge (FG) and one cluster (MTG) was located outside the MD. Thus a small number of regions processing visual body or face information present in the static frame, increased their response to this period, possibly as a trigger for neighboring Early- and Entire-VE leads.

**Table 3:** characteristics of the SE clusters.

| RegionName | # of patients | # of leads | AvgMD |
| --- | --- | --- | --- |
| FG | 2 | 6 | 0,53 |
| IFS | 2 | 3 | 1,68 |
| IPS | 1 | 2 | 4,88 |
| LO | 4 | 7 | 1,50 |
| MTG | 2 | 4 | -0,49 |

### Properties of Early-VE Clusters

The five Early-VE clusters were located in the dorsal occipital cortex (LO cluster), along the IPS (IPS cluster), in the dorsal bank of the POS (POS cluster), in the OTS (OTS cluster) and in the posterior parts of MTG and STS, referred to as posterior Temporal (pTC) cluster (fig 3, S4). The time courses of z-scored gamma power in the four conditions (fig 5) showed that activity in all five clusters increased in the short action trials during the first 500ms of the dynamic period, starting in IPS, pTC, and to some degree, POS clusters already during the static period. Such increases in short trials were not observed in the gender discrimination task, indicating the task specificity of these increases. The long trials (blue) showed further task dependencies: the IPS, OTS, and LO, which all belong to the MD system, increased their activity in response to the static onset which was actually stronger in the gender task. But in the long action trials these clusters showed also a response to the video, more clearly in the OTS and IPS clusters. The pTC cluster showed a weak response to static frame and the video, with little difference between tasks, while the POS cluster showed little response at all in long trials.

**Fig 5:**
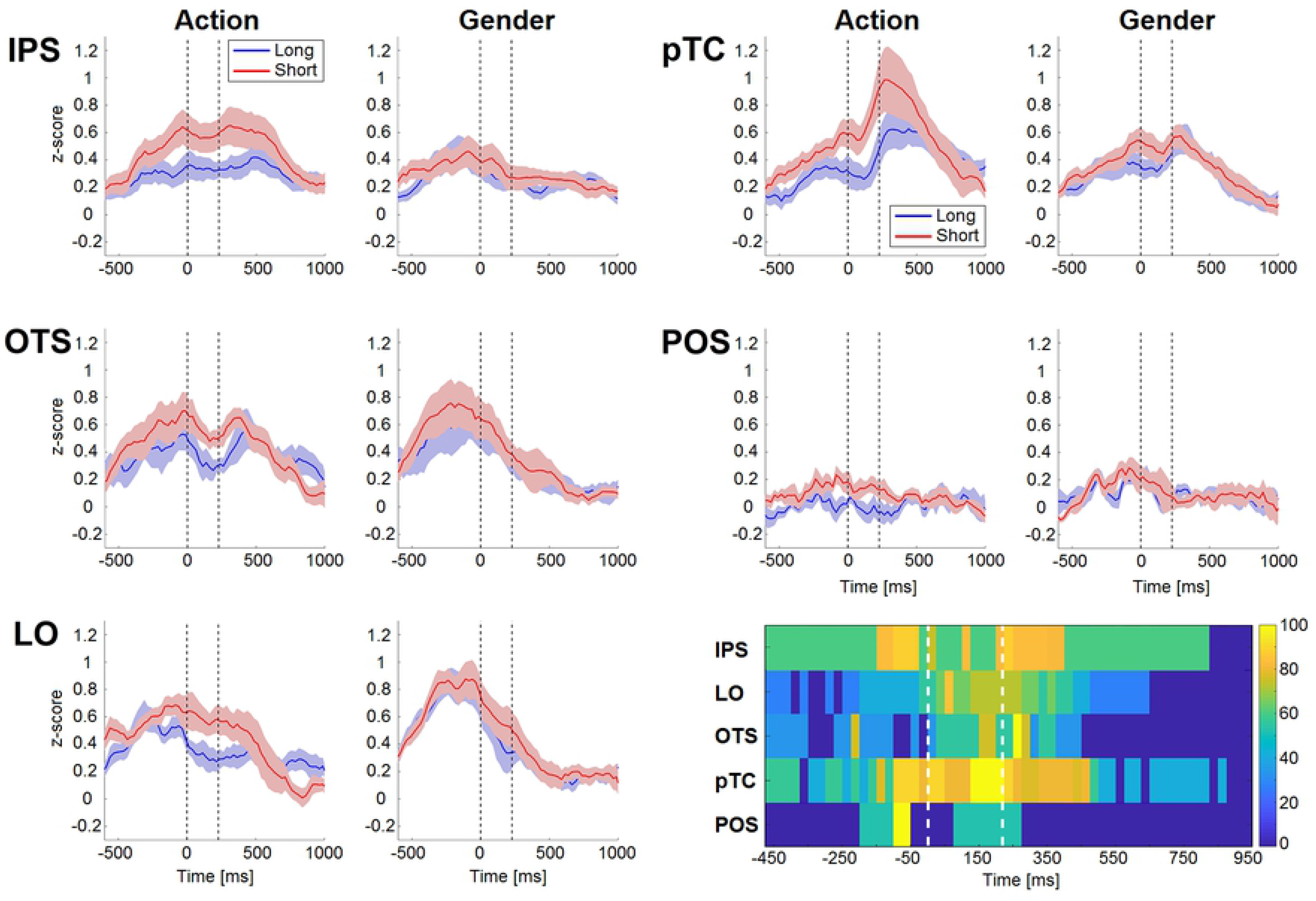
Average time courses of the 5 Early-VE clusters in the four conditions (red short, blue long); left and right clusters: inside and outside MD system respectively (see fig S4). Bottom right significance plots for the five clusters: color code: average (over patients) proportion of leads showing a significant increase (sliding window t test). Same conventions as fig 3.

The average curves (fig 5) show the variability across patients, providing coarse estimates of the periods during which activity significantly increased in the short action trials. The significance plots (see methods) provide the precise timing of the activity increases in short action trials for the different Early-VE clusters, using a color code for the mean proportion of significant leads (lower right panel of fig 5). The increase started early in the IPS cluster, reaching near maximum values from 150ms before until 450ms after video onset. The increase occurred much earlier in the IPS cluster than the two other MD clusters, even if in the basic task information (about observed actions) flows in the opposite direction, according to anatomical data in the monkey (Nelissen et al 2011). Short-trial increases in pTC and POS (both outside the MD) started about simultaneously the IPS, the increase in POS being very short-lived.

The time course of the early-VE clusters, and their location in regions known to belong to the action observation network (Cross et al 2009, Jastorff et al 2010), suggest that these clusters are the effectors of the effort response, enhancing, as expected from the behavior (see above), the visual representations of the observed manipulative actions. This predicts that their activity should correlate positively with accuracy and negatively with RT. We computed for each lead the difference in z-scored gamma power between correct and error trials, averaging over a window 50-550ms after video onset, matching the timing of Early-VE leads increases (fig 3). The distribution of these differences (fig 6) was significantly larger than zero, both for all 64 Early-VE leads (two tailed t test, t=2.37, df 63, p<0.025), and for the 20 leads in the MD clusters (two tailed t test, t=2.67, df 19, p<0.02). For comparison, the distribution of these differences for Late-VE leads was close to zero (mean value 0.03). The distribution of the correlation between activity and RT was skewed to negative values as well for all 64 Early-VE leads as the 20 leads in the MD clusters, but this was non-significant, perhaps due to the small range of RTs. Indeed, when plotting the activity as a function of RT across leads, the negative correlation reached significance (r= −0.51, p<0.02) for error trials, covering a large range of RTs, but not correct trials (r= −0.27, p>0.2) with a smaller RT range.

**Fig 6:**
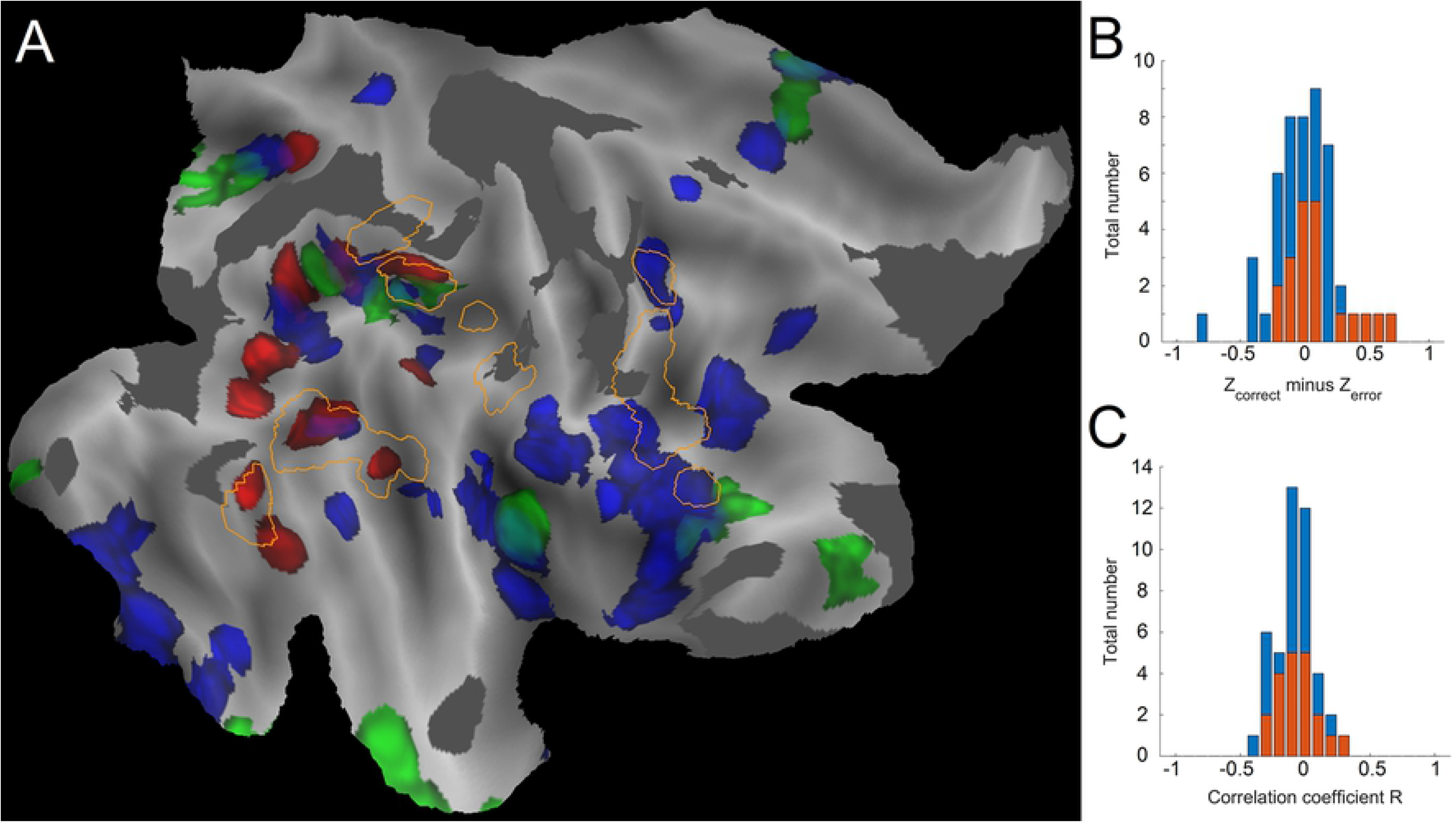
Properties of Early-VE leads; A: flatmap of right hemisphere showing the 31 VE clusters (color coded as in fig 3) with the mask of action observation taken from Caspers et al 2010 (orange outlines), B distribution of the difference in z-score for correct and error trials for the 64 Early VE leads and those in the MD clusters (orange); C: distribution of r values of correlation between RT and z-scored gamma in the 64 Early VE leads and those in the MD clusters (orange).

To further support the role of the Early-VE clusters as effectors of the effort reaction, we compared their location to the map of action observation regions obtained by a meta-analysis (Caspers et al (2010). This map overlapped, as expected, mainly with Early-VE clusters (fig 6), those in lateral occipito-temporal cortex (LOTC) and in rostral IPS. In addition, it overlapped only with the mPreCG Entire-VE cluster, and the aIPS Entire or Late-VE clusters, neighboring the rostral part of the IPS Early-VE cluster (fig 6). Comparing figures 6 and 3 shows that the action observation regions overlap to some extend with the MD system, especially at the level of the IPS. This explains that several Early-VE clusters, although effectors of the effort response, were located in the MD system, supposedly controlling the effector regions. Figure S6 details the overlap between the VE clusters and action observation map in rostral IPS. The dorsal intraparietal sulcus anterior (DIPSA) and putative human anterior intraparietal sulcus (phAIP) ellipses (Orban 2016), both action-observation regions, overlap with the rostral IPS Early-VE cluster, and with aIPS Entire and Late-VE clusters, respectively. These ellipses, homologues of the posterior and anterior parts of monkey AIP (Orban 2016), which hosts OMA selective neurons (Lanzillotto 2019), have been implicated in the discrimination of observed manipulative actions (Orban et al 2019). Hence, we computed the average time courses of the activity in these ellipses for all four conditions (fig S10). The DIPSA leads responded to the static presentation in both tasks, accompanied in action discrimination by a response to the video, larger in short than in long trials. In the phAIP ellipse (fig S6) the difference between short and long action trials was more striking, with an opposite task dependence for static and video responses. In short gender trials little changed, but in the short action trials both the activity during static and dynamic period were enhanced. Thus global activity in these ellipses was task dependent, being stronger in the beginning of the video period in short than long action trials. Hence, these ellipses, dominated by the Early-VE pattern but including also Entire- and Late-VE leads, may represent the effector regions which interact with the controlling MD system.

### Properties of Entire-VE clusters

The majority of the VE clusters belonged to the Entire type. Twelve of them, all located in prefrontal or parietal cortex, belonged to the MD system, while a further 6 clusters in temporal cortex or at the edges of the parietal cortex were located outside the MD system (fig 3, S4). Their time courses (fig 7 and S7 for those inside the MD system, fig S8 for those outside) confirm that the increase in the short trials extended beyond the first 500ms of the dynamic period, finishing at variable times before the response. In some clusters, the increase also involved the last part of the static period. In all 18 clusters the increase in the short trials was task specific, being absent in the gender task (fig S7, S8), with the exception of the IFS cluster showing a weak, late increase in short gender trials. Contrary to the Early-VE clusters, most Entire-VE clusters showed little response in the long action trials, with the exception of video period in IFS and pMTG, and static period in mIPS (fig 7, S8). Finally, it is worth pointing out that the 6 Entire-VE clusters outside the MD system were again an *unpredicted finding* (labeled 3 in fig 2), as the Entire-VE time course was expected for control regions. Three of the clusters outside the MD system (fig S8) showed a response in the long gender trials: clearly in TO, but also OP1-4 and pSTS, suggesting that they process features related to the actor, rather than the action.

**Fig 7:**
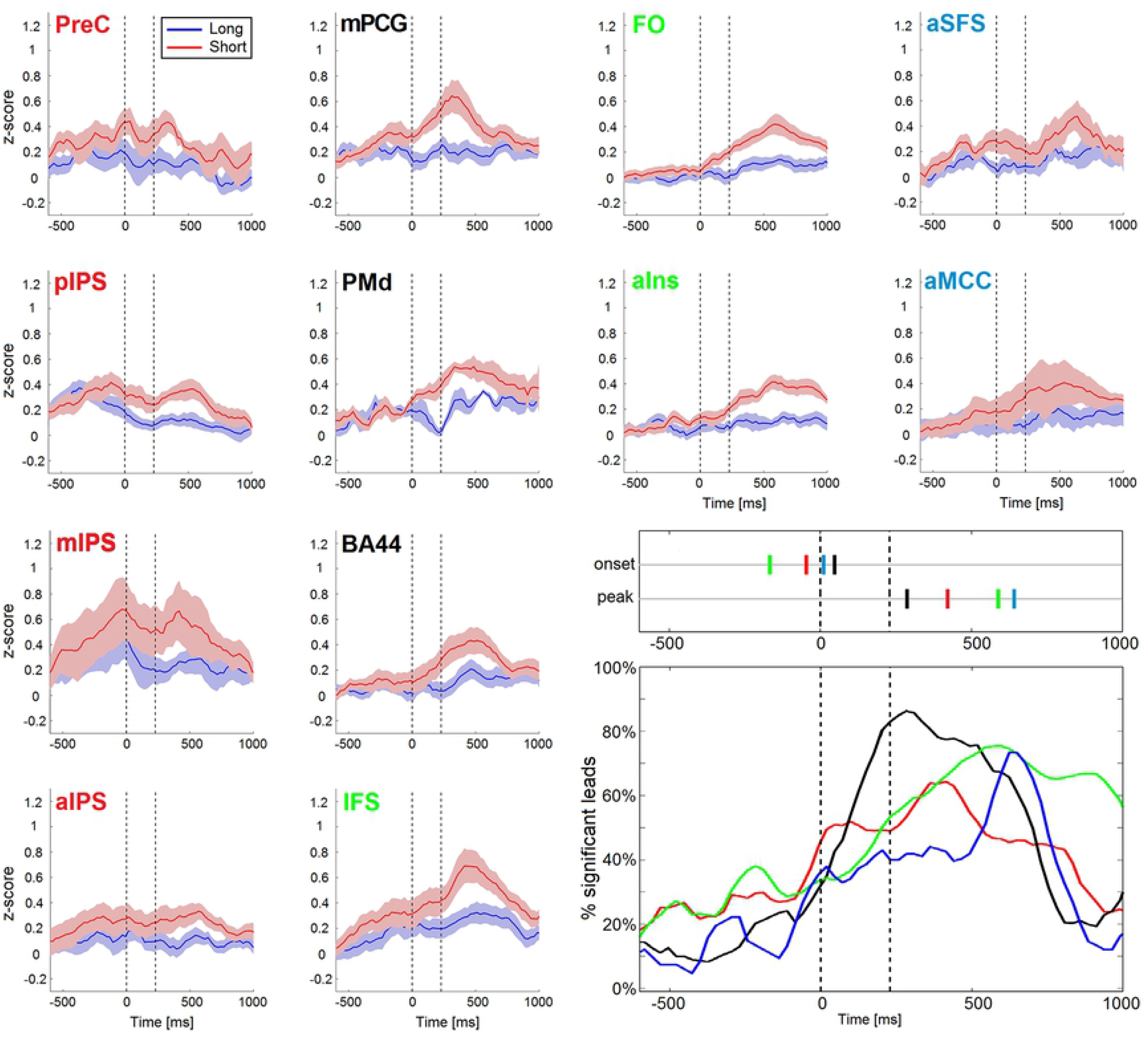
Average time courses of the 12 clusters of Entire-VE leads (see fig S4) inside the MD system in action trials (red short, blue long), grouped into four anatomical regions: PPC, cPFC, vlPFC, and dPFC. Bottom right significance curves for the four anatomical groups (color coded). Vertical lines above the significance curves indicate onset latency (50% of maximum) and peak of the curves. For gender trials of these 12 clusters see fig S7. For Entire-VE clusters outside the MD system see fig S8.

To clarify the dynamics of the increases in the 12 Entire-VE clusters of the MD system we grouped them anatomically: four clusters (3 along the IPS and one in the precuneus) in posterior parietal cortex (PPC), three (mPreCG, PMd and BA44) in caudal PFC (cPFC), another three (IFS, FO and aIns) in ventro-lateral PFC (vlPFC), including the anterior insula, and two (aMCC, aSFS) in dorsal PFC (dPFC). We computed *significance curves*, plotting grand averages of the proportions of significant leads per patient over time, for these four groups (fig 7 bottom right). The percentage of significant leads was slightly larger in the 3 PFC regions than in PPC, and the curves reached their maxima: first in cPFC, closely followed by PPC, then vlPFC and finally dPFC. However, as expected for Entire-VE clusters the increases reached already some level of significance earlier on, captured by setting 50% of the maximum of the curve as threshold defining onset latency (Siegel et al (2015), vlPFC regions showed a gradual increase in the proportion of significant leads, reaching transiently 50% of the maximum well before video onset. The other 3 regions had clear-cut onset latencies: PPC slightly before video onset, dPFC just after, and finally cPFC, about 50ms after video onset (fig 7). Significance rose more rapidly for this latter group than the others. Thus the effort-related activity increases in the MD system did not follow a simple sequential pattern, but a complex one where regions starting early reached their maximum much later.

Given their time course and localization, the Entire-VE clusters are likely to exert the increased control in the difficult trials. The hallmark of control being error monitoring, we compared the responses in short action trails for correct and error trials. To guarantee sufficient error trials, we considered only patients reaching at most 91% accuracy in short action trials, and in those patients only leads with at least 5 error trials. This left at least two patients and two leads in all 12 Entire-VE clusters inside the MD system, with a median of 5 patients and 7 leads per cluster. For each cluster, we compared the activity in correct and error short action trials using a sliding t test as before, to provide the timing of error-correct differences. These differences reached significance in a sizeable proportion of leads in five of the 12 clusters, but with two different time courses. To enhance representability we grouped clusters this time by effect and location, yielding four functional groups (fig 8). The earliest error signals appeared in parietal regions mIPS and Prec (fig 8A,E), reaching maximum significance at 450ms after video onset, and again around 950ms after video onset. Error signals in two caudal PFC clusters (mPrecCG and BA44) and aMCC (fig 8B,F) reached maximum significance at 800ms after video onset. Another group of PFC clusters (IFS, FO, aIns) showed only marginal increase at the end of the dynamic epoch (fig 8C, G), while a final group of four clusters (pIPS and aIPS in PPC, and PMd and aSFS in PFC) showed no effect (fig 8D, H). In fact, the two PFC clusters of this latter group, showed an opposite effect (more activity for correct than error trials, fig 8D), first in PMd, and later in aSFS. It is worth stressing that these error signals were specific to the Entire VE clusters (fig S9), and occurred before the response, thus predicting errors rather than monitoring them (Tang et al 2016).

**Fig 8:**
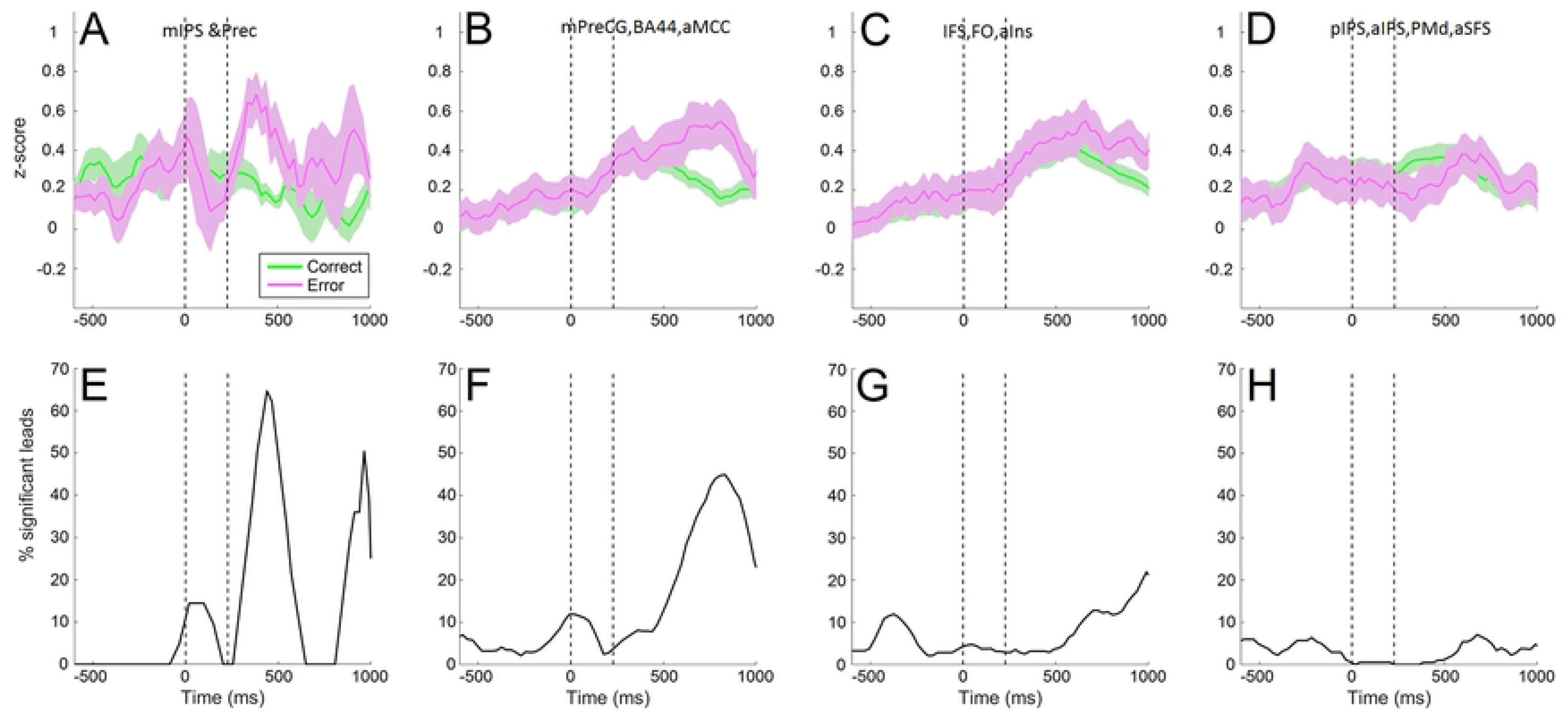
Comparison of error and correct trials for short action trials in four groups of Entire-VE MD clusters: mIPS & Prec (A, E), mPReCG, BA44 & aMCC (B, F), IFS, FO & aIns (C, G) and pIPS, aIPS, PMd & aSFS (D, H). A-D: average time course of activity in short trials for error (light purple) and correct (green) trials, E-H: statistical significance: average (across patients) proportion of leads showing a statistical significance (sliding window t test). For similar curves for the Early-VE and Late-VE clusters see fig S9.

Again, the time courses of error prediction was complex, starting in PPC, spreading to PFC and returning to PPC.

### Properties of clusters with unexpected time courses

While we expected increases in the short action trials, either at the beginning or the over the entire dynamic period, we also observed clusters of leads increasing their activity near the end of the dynamic period and during the static period.

Five VE clusters, all inside the MD system, showed a Late-VE pattern. They were located in parietal cortex (IPS and POS), and PFC. As expected the increase in short action trials started after 500ms into the dynamic period (fig 9). The time-courses confirmed the distinction between Entire-VE and Late-VE clusters with similar location (eg in pIPS or aMCC). The increases were slightly less task specific, as two clusters showed a weak increase in the short gender trials (fig S10). Generally these clusters showed little response during the long action trials, with the exception of pIPS.

**Fig 9:**
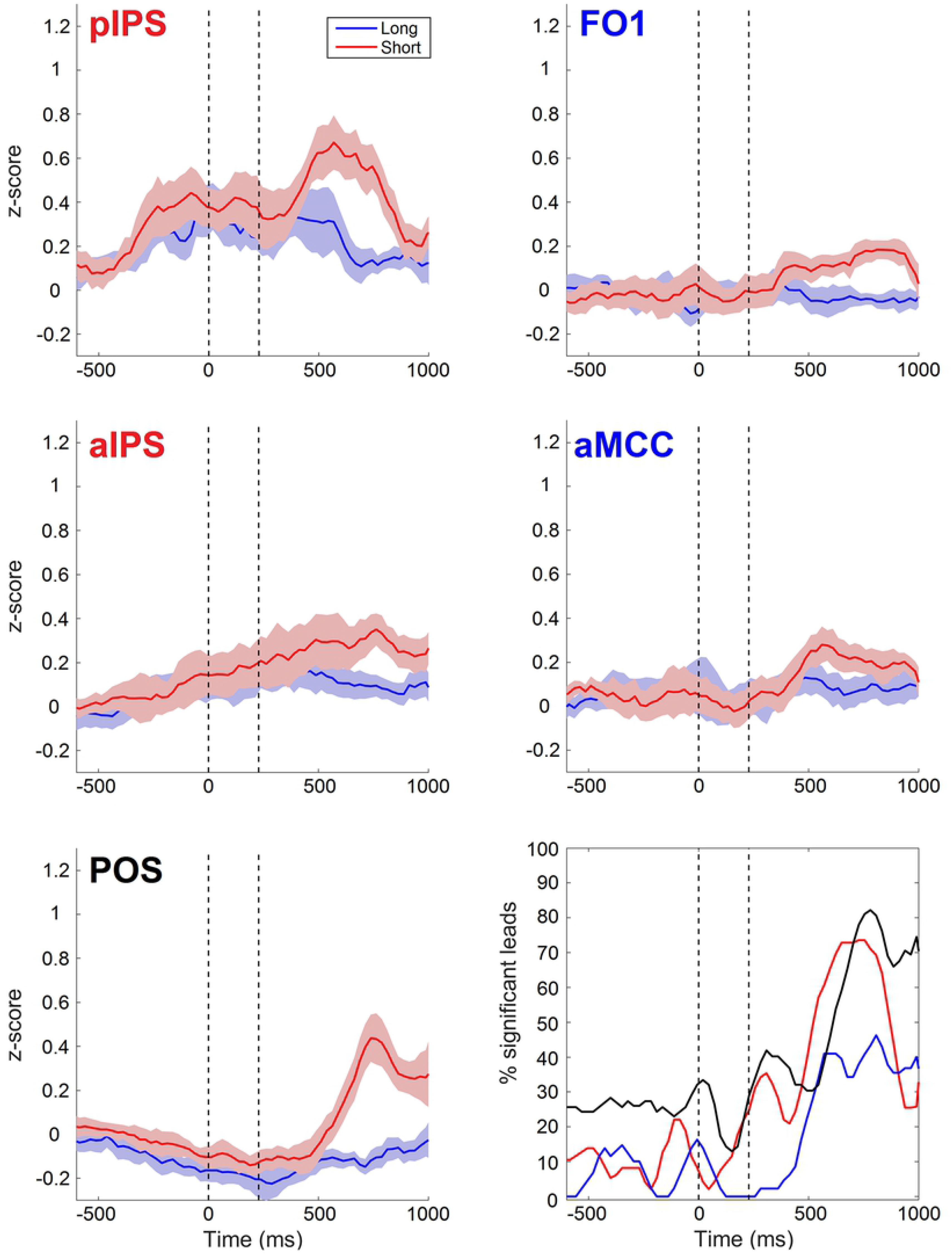
Average time courses of the 5 clusters of Late-VE leads (see fig S4) inside the MD system in action trials (red short, blue long), grouped into 3 regions (IPS, PFC and POS). Bottom right significance curves for the three regions (color coded). For gender trials of these 5 clusters see, fig S10.

Again we grouped the 5 Late-VE clusters by anatomical location, and computed significance curves (fig 9). Using the same criterion for onset of significance as used for the Entire-VE groups, significant increase (ie 50% of maximum) started at about 460ms after video onset, well after the Entire-VE groups, which all started before or around video onset. Onset was very similar for the IPS and PFC clusters, but the IPS clusters reached their maximum slightly earlier (700ms after video onset) than the PFC (780ms after video onset). The curve of the POS cluster showed an ambiguous onset between 270 and 540ms after video onset, but reached its maximum as late as the PFC Late-VE clusters (780ms).

Four of the SE clusters (the MTG cluster being the exception) were located near but outside the action observation map (Caspers et al 2010), as expected since the meta-analysis generally included static controls. Importantly for the presumed function, the LO, MTG, and IPS -SE clusters had one lead in common with the LO, pTC, and IPS Early-VE clusters respectively, and the IFS-SE cluster even had two leads in common with its Entire-VE counterpart, explaining its early onset.

The time courses of the increased static activity in the short compared to long action trials and the significance plots (fig 10) clearly set the IPS SE cluster apart: the increase started earlier (despite a longer latency in long action trials), and lasted longer than in the other clusters. In most clusters the increase in short trials was specific for action trials, except in the MTG part of EBA. These results support the view that these SE leads, in particular those in IPS, serve as a trigger for the increase in Early and Entire VE clusters.

**Fig 10:**
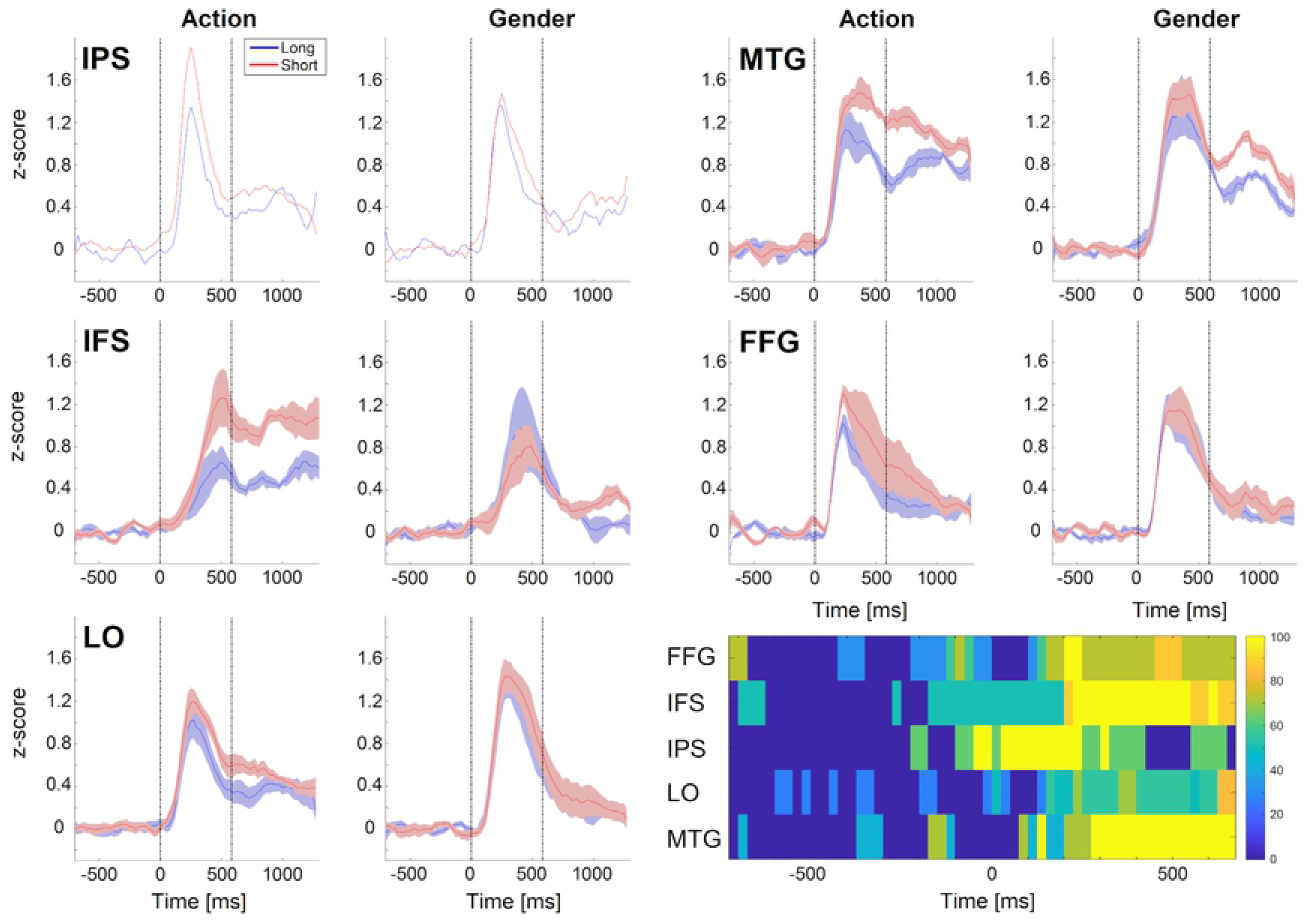
Average time courses of the 5 clusters of static enhanced leads in the four conditions (red short, blue long); bottom right; significance plots of the five clusters. Vertical lines indicate average static period.

## Discussion

Our results show that when subjects face a difficult action discrimination they indeed call upon the MD system, of which we documented both the spatial and temporal complexity. They also illuminate the way the MD system interacts with other brain systems, and the role of the PPC in action observation.

### Comparison with earlier studies

In the present study, most regions of the MD system (Fedorenko et al 2013) increased their activity when the patients performed a difficult action discrimination. This increase was specific to the difficult task, and did not simply reflect the shortening of the videos. The short period during which visual information about actions was provided, may explain that we failed to observe the increase in RT, which typically accompanies the decrease in accuracy in difficult tasks (Lingnau & Petris 2013). The localization of the entire-VE clusters followed the spatial distribution of the MD system closely (fig 3). Yet, in the present study the difficulty arose from the truncation of the videos, of which the duration was reduced to 228ms on average. Hence, our results fail to support the earlier view (Han & Marois 2013, Wen et al 2018) that the MD system does not cope with difficulty arising from data limitation. In our experiment, the single difficult condition increased the error rate by 15%, considerable compared to error rates of a few percent induced by conflict (Tang et al 2016, Smith et al 2019), but less than the 30% in Wen et al (2018) or the 25 & 15% in Han & Marois (2013) experiments. However, these two studies also differed in the recording technique, fMRI being less sensitive, especially for short-lived activations, than stereo-EEG. This factor is likely more important, as it fits the pattern of differences with Han and Marois (2013), who obtained fronto-parietal activation when they introduced a distractor, but not when they reduced the duration of the stimuli. Thus whether and why the MD system fails to cope with difficulty in simple sensorial discriminations needs further work, taking into account the sensitivity of the technique used.

It has been long recognized that control is costly (Botvinick & Braver 2015). Three types of costs have been considered in effortful tasks (Shenhav at al 2017): 1) metabolic depletion, 2) computational limitations in the capacity for controlled processing, and 3) restrictions in control-dependent behavior reflecting cross-talk arising from local bottlenecks in processing, when different tasks compete to use the same set of representations or apparatus for different purposes. The present experiment using a single task involving a single item, provides strong support for metabolic costs playing an important role in effortful tasks. This consideration may also account for the lack of tonic effects during baseline in the difficult sub-blocks, a result in agreement with those of Dubis et al (2016).

### Complexity of the MD system

The present results illustrate dramatically the complexity of the MD system which includes no less than 17 distinct VE-clusters, excluding the 3 Early-VE clusters overlapping with the MD system. The latter overlap indicates that the MD system cannot be defined purely on anatomical grounds, but requires the combination of dynamics, function, and localization (used to exclude the 3 Early-VE clusters, see below). The 17 MD clusters were located along the IPS but also on the edges of the PPC in the precuneus and the parieto-occipital cortex, as well as in various parts of the lateral PFC and aMCC. Importantly, the excellent fixation performance of the patients eliminate the possibility that the enhanced activity in IPS in short action trials reflected increased oculomotor activity.

The majority of the MD clusters (12/17) are active throughout the entire dynamic period (Entire-VE-clusters). In fact, they started to increase their activity early on, close the video onset, but reached their maximum in the second part of the dynamic period (Table 4). We take the variety of time courses as a suggestion that these Entire-VE clusters may play different executive roles, and propose the following tentative scheme. The vlPFC clusters start the process gradually, in keeping with the overlap of the IFS VE cluster with a SE cluster. This gradual onset is thought to represent an increased task-set (Dosenbach et al 2006, Higo et al 2011), ensuring the task is performed despite the difficulty, which is aversive (Shenhav et al 2017). In addition, at this early stage, the anterior Insula may mark the static frame and the ensuing brief video as salient events (Menon and Uddin 2010), to improve their processing. Next, the dPFC region increases its activity, which, we suggest, represents the decision to increase control (Shenhav et al 2017). This signal is then boosts the activity of cPFC, which we interpret as ensuring that control is enhanced, followed by a peak in PPC activity. Thus in the MD system, the PPC follows the PFC, and likely provides the link with the effector regions. As the dynamic period ends, vlPFC reaches its maximum, which may represent, the evaluation of the costs of control (Shenhav et al (2017), mainly metabolic in nature (see above), in line with the implication of neighboring anterior insula, receiving signals related to hypoglycemia, hypoxia or metabolites from muscles through a direct thalamic pathway (Craig 2003). Finally, dPFC reaches it maximum, which, we propose, integrates costs and benefits (Shenhav et al 2017), the latter represented by correct performance enhancing activity in aSFS.

**Table 4:**
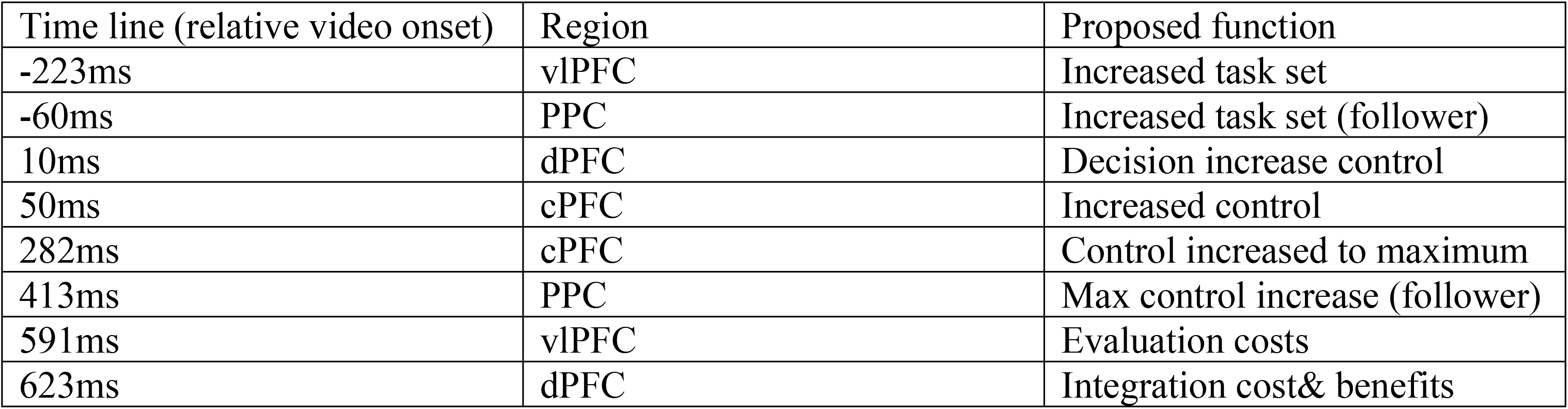
time line of the increased activity in Entire-VE cluster groups.

Five of the 17 VE clusters in the MD system increased their activity only in the second half of the dynamic period (Late-VE clusters). Since the increase occurs so late, too late to be used in control itself, we propose that the Late-VE clusters perform an evaluation function within the MD system. Specifically, the aMCC is thought to be the hub of these Late clusters, continuing the integration started by its Entire-VE counterpart (Table 4), to address two questions: 1) was the strategy to cope with the difficulty adequate, and 2) was the effort worthwhile? Answering these questions may rely on signals from parietal and orbitofrontal cortex. Such evaluation may be critical to improve later efforts and account for the contribution of the MD system to fluid intelligence (Tschenscher et al 2017). The role of the POS late-VE cluster, with its complex time course, is less clear. POS might correspond to area prostriata (Mikellidou et al 2017), providing low-level visual signals directly to the aMCC (Morecraft et al 2012) possibly to inform the subject about his own situation in the environment.

One important finding of the stereo-EEG recordings is the predictive nature of the error signals handled by the MD system. These signals arose in PPC shortly after the end of the truncated video and travelled back and forth to the cPFC, and aMCC, the role of which in error correction is well known (Carter et al 1998). These error signals occurred in the MD system before the subject actually made mistakes, this predictive nature being computationally advantageous as it allows feed forward control. We propose that these error-predicting signals in fact, reflect the weakness of the sensory evidence, as proposed by Kiani and Shadlen (2009) for confidence processing in monkey LIP. Indeed, similar signals have been recently recorded (Desenter et al 2019) over human parietal cortex, where we found the error predictions to start, and they have been postulated to underlie the so-called sensory confidence of decisions (Gherman & Philiastides 2015). The single cell recordings (Kiani and Shadlen 2009) suggest a mechanistic account for the generation of error predictions. If some time after video onset in the short trials the decision variable remains too close to zero, an error could be predicted. The OFF responses generated by the end of the video could signal the time at which the variable needs to be evaluated, explaining the appearance of error prediction signals in PPC shortly after the end of the video.

### Action observation circuits

The MD system supposedly (Fedorenko et al 2013) interacts with the regions involved in the basic task, to enhance the processing in these regions, for the task to be completed successfully (see Power & Petersen 2013 for a similar distinction between control and processor regions). In the present case, we expect the MD system to interact with the action discrimination circuit. Psychophysical results (Platonov and Orban 2016) suggest that such circuit, as many others (Palmer et al 2005), includes an initial specific sensory step, collecting the evidence about observed actions and a second decision step accumulating the evidence. Given the shorter RT and lower accuracy of short action trials, the activity in the latter step is likely to be unchanged or even decreased during the short trials (see above). Thus the MD system should interact with the specific sensory regions of the action discrimination circuit (fig 2), to which the early-VE clusters correspond, even if there is some overlap with the MD system. Indeed Early-VE leads are active during the long action discrimination trials, while the Entire-and Late-VE leads are not (fig 3) This distinction holds at the individual cluster level: most Entire-and Late-VE clusters do not respond during long action trials (fig 5,6), while at least three Early-VE clusters do: IPS, OTS and pTC (fig 4). Since these 3 clusters also overlap with the .AON (fig 8), engaged in passive viewing of observed actions, we propose these three regions as the candidate effector regions of the difficult action discrimination task. The positive correlation of the Early-VE activity with accuracy (fig 6) and the weak negative correlation with RTs support this identification as effector regions. Amongst the regions overlapping with the action observation network (Cross et al 2009, Caspers et al 2010, Jastorff et al 2011), IPS stands out as starting the increased processing in the short action trials, and in particular DIPSA and phAIP in rostral IPS. Both regions show, on average, an increase in the short compared to long action trials during the end of the static period and the short video (fig S6). The increase in short action trials was slightly stronger in phAIP than in DIPSA, in keeping with its stronger video response in the long action trials. phAIP, including also Entire-VE and Late-VE leads, is a candidate region for linking the action discrimination circuit with the MD system.

These results also show that action-observation regions in LOTC and parietal cortex remained clearly active during the short action trials, consistent with the results of Tucciarelli et al (2015). These authors decoded observed action (833ms long videos) with 52% accuracy from low frequency (<25 Hz) bands in MEG earlier in LOTC and left superior parietal lobule (SPL) than in premotor cortex (720ms after action onset). They suggested that the similarity of the onset latency and the duration threshold (measured separately) favored the LOTC as a critical region for action observation. The present results point to the weakness of such an inference, as reducing video duration does much more than cutting off responses, and latency may change with video duration. Combined with earlier studies they support the role of PPC in action observation. phAIP was more activated in long action than gender discrimination, in agreement with a recent fMRI study (Orban et al 2019). Furthermore this region, and its homologue in the monkey, AIP, has been shown to house observed manipulative-action selective neurons (Lanzilotto et al 2019, Aflalo et al 2018), well suited to represent observed action identity. Such selectivity suggests an increase in the gain of the selectivity curves as a mechanism for boosting sensory evidence about observed actions.

### Interactions of the MD system with other cortical systems (fig 2)

We expected increased activity during the difficult action discrimination trials in the MD system and the action discrimination circuit, respectively control and effector regions. As discussed above, these predictions were born, but our results revealed two more sets of enhanced cortical regions. One such system were the five SE clusters, responsive to the specific static information present in the frame shown before the truncated video, a possible trigger signal for the VE clusters. These clusters correspond to regions known to process the human body and face: EBA (Downing et al 2001, Moro et al 2008, Weiner & Grill Spector 2013, FBA (Peelen & Downing 2005) and FFA (Kanwisher et al 1997), the prefrontal face region (Tsao et al 2008) and a region in IPS dorsal of that processing views of limbs (Orlov et al 2010). EBA included LO and MTG parts, as expected from the description of EBA surrounding the MT cluster (Weiner & Grill-Spector 2013), but the functional properties of these two parts (fig 10) were clearly different. The SE-clusters are distinct from the Early-VE clusters, except for some overlap at early processing levels (LO and pMTG). Indeed Downing et al (2006) has shown that EBA contributed to action perception by processing the static structure of the human form, the deformation of which is one the signals used to extract observed actions from the retinal array (Van Geneugden et al 2009). Although distinct, the SE clusters had a few leads in common with VE clusters: the IPS SE cluster with the IPS Early-VE cluster and IFS with the IFS Entire VE cluster. The IFS SE-cluster, although starting later than the IPS SE cluster, may be the main trigger given its strong lasting effect. Thus to avoid tonic signals (Summerfield & Lange 2014), which are metabolically costly, the MD system may have behaved opportunistically and recruited a small set of cortical regions, unrelated to the action task, to serve as a warning system for the occurrence of a brief video.

Our results revealed a second unexpected set of enhanced regions: several of the Entire-VE clusters located outside the MD system, processing information present in the video other than action identity, to bias choices of the observed action. One cluster, located in TO and neighboring STG, corresponds to regions shown to represent the speaker (Belin et al 2000) and more generally the actor as a human communicating by voice and body language (Corbo and Orban 2017). Its role as processing actor information is supported by its activation in the gender task, with a time course clearly different from the fast activation in ATL, known to underlie the gender discrimination (Platonov et al 2019). Two regions exhibiting similar activity in the gender task, may play an analogous role: one region includes the pMTG and pSTS clusters, located near regions processing the actor as an agent behaving rationally (Jastorff et al 2011), the other are the opercular regions OP1-4. SII neurons in the monkey has been shown to respond to observed actions (Hihara et al 2015) and these visual signals may, we speculate, process visual information about the texture of the actor skin, and hence indirectly his/her physiological state. In addition to the actor, the video also contains information about the scene, which might be processed by the parahippocampal Entire-VE cluster (Epstein &Kanwisher 1998). Finally, the location of the POS Entire-VE cluster suggests a possible homology with V6A, the neurons of which process object-related information, when the object is handled by another actor (Breviglieri et al 2019). Thus these Entire-VE clusters outside the MD system may provide object, scene and actor-related information to bias the choice of observed action, a process that may already start in the neighboring POS and pTC Early-VE clusters (also outside the MD system). These results document another property of the MD system which searches for additional, even weak, information to increase chances of successfully solving a difficult task, as if it was ‘pondering’ this weak information.

### Limitations of the study

Although we recorded from over 3000 cortical sites in 28 patients, we sampled the right hemisphere much more than the left. As we did not find obvious asymmetries in the different types of enhanced leads, and the MD system is completely bilateral (Fedorenko e al 2013), the detailed description of the right hemisphere may be valid for both hemispheres. But even the right hemisphere was not sampled evenly, such that the lack of Early-VE leads in the premotor cortex may reflect under-sampling of that region. Yet the VE clusters matched the extent of the MD system rather well, indicating that sampling, while uneven, was globally adequate.

To provide more representative results we concentrated on clustered VE leads, using only seventy percent or less of the VE leads in our functional descriptions. Combined with the previous limitation, it is conceivable that more clusters will be detected when sampling the hemisphere more densely. This second limitation is mitigated by the observation that leads with strong MD index (t>1.5) were far less common amongst isolated (14%) than clustered (38%) VE leads, especially Entire-VE leads involved in control. Thus, by concentrating on clustered leads we may have focused on the MD system.

Finally, these invasive recordings were made in drug resistant epileptic patients. The inclusion criteria were very strict and included absent or very limited anatomical anomalies. Most patients scored normally on the neuropsychological tests, all of them fixated very well during the tasks, and most importantly, they performed the action discrimination as well as normal, young test subjects (Platonov & Orban 2016), both for long and short trials. The cortical leads were carefully inspected for IEDs, and those trials presenting IEDs, generally a small number (see Platonov et al 2019), were removed from analysis. Furthermore, IEDs are unrelated to the stimulus or the response of the subject and therefore contribute little to the time courses synchronized to these external events. In the right hemisphere only 29 out of 432 VE leads (7%) and none of the 39 SE leads were located in the EZ. Only 2 Late-VE clusters (ATL and aFO), both outside the MD, included more than half the leads from the EZ and were excluded from the analysis. Of the remaining 28 clusters, 17 included no leads in the EZ, 6 included few (20% or less) EZ leads, and only five included 30% leads from the EZ; The two of these, belonging to the MD system, were shown to behave similarly, whether the EZ leads were included or not. Therefore, the functional properties of the leads documented in the present study represent physiological properties of the human brain, measured with a sensitivity and temporal resolution offered by no other method of investigation.

### Conclusions

Our results prove the involvement of the MD system in difficult action discrimination, thereby revealing novel features of the MD system: its complex dynamics, but also the search for additional information, the evaluation of its own functioning, and the predictions of errors. It underscores the strength of stereo-EEG, the sensitivity and excellent spatio-temporal resolution of which has enriched our understanding of the MD system.

## Material and Methods

### Patients

Stereotactic Electroencephalography (stereo-EEG) data were collected from 28 patients (13 female, age 18-49, mean 32 years, Tables S1&2) suffering from drug-resistant focal epilepsy. Intracerebral electrodes were stereotactically implanted at the Claudio Munari Centre of Epilepsy Surgery, as part of the presurgical evaluation to define the cerebral structures involved in seizure generation (Munari et al 1994, Cardinale et al 2019). The implantation was based on the presumptive location of the epileptogenic zone (EZ), derived from clinical history, examination of non-invasive long-term video-EEG monitoring, and neuroimaging. Patients were fully informed about the electrode implantation and stereo-EEG recordings. The present study received the approval of the Ethics Committee of Niguarda hospital (ID 939-2.12.2013) and informed consent was obtained from all patients in the study. Data of 24 of the patients, performing the same tasks, were analyzed for a different purpose in Platonov et al (2019).

#### Inclusion criteria

As described in Platonov et al (2019), patient selection was based on a series of stringent anatomical, neurophysiological, neurological and neuropsychological criteria, to minimize the recording of any data from functionally compromised sectors of the brain tissues. Only patients whose magnetic resonance imaging (MRI) showed no anatomical abnormalities, or only very restricted anomalies (e.g. focal cortical dysplasia) were included in the study. Neurophysiological examination included the inspection of the EEG tracks at rest, during wakefulness and sleep from both intracranial and scalp EEG. Pathological activity was characterized by the presence of epileptic discharge at the seizure onset, but epileptic spikes could be present in leads exploring the regions surrounding the EZ during the interictal periods. Any trial presenting interictal epileptic discharges (IEDs) at any latency during the stimulus presentation was removed. In addition to the spontaneous EEG activity, the neurophysiological investigation also included an assessment of the normal reactivity of both intracranial and scalp EEG to a large set of peripheral stimulations (somatosensory, visual, vestibular and auditory stimulations) to verify normal conduction times and overall sensory reactivity. On the neurological side no seizure, no alteration in the sleep/wake cycle, and no change in pharmacological treatment should have taken place within the last 24h before the experimental recording of a patient included in the study. Neurological examination had to be unremarkable, with in particular absence of motor or visual deficits. Finally, a series of neuropsychological tests was administered by experienced neuropsychologists (see supplementary info).

#### Localization of recording sites with respect to lesions and epileptogenic zone

Only patients presenting either no anatomical alterations (n = 20) or only small abnormalities (n = 8), as evident on MRI, were included. Five of the 8 patients with positive MRI, showed minimal periventricular nodular heterotopia (4 in the temporal lobe, one in the occipital lobe), two patients focal cortical dysplasia (FCD) in the frontal lobe, and one hippocampal sclerosis.

The epileptogenic zone (EZ) involved generally parts of temporal or frontal cortex. Its extent was established in each patient and the overlap with clusters of leads showing significantly increased activity with difficulty determined.

### Electrode Implantation

Most implantations were unilateral, because clinical evidence generally indicates the hemisphere generating the seizures. Only 3 of the 28 patients were implanted bilaterally, resulting in a total of 31 implanted hemispheres. An average of 16 depth electrodes (range: 12–21) were implanted in different regions of the hemisphere using stereotactic coordinates. Each cylindrical electrode had a diameter of 0.8 mm and consisted of eight to eighteen 2-mm-long contacts (leads), spaced 1.5 mm apart (DIXI Medical, Besancon, France). Immediately after the implantation, cone-beam computed tomography was obtained with the O-arm scanner (Medtronic) and registered to preimplantation 3D T1-weighted MR images (Avanzini et al 2016). Subsequently, multimodal views were constructed using the 3D Slicer software package (Fedorov et al 2012), and the exact position in the brain of all leads implanted in a single patient was determined by using multiplanar reconstructions (Dale et al 1999).

### Behavioral testing

Behavioral testing of the patients was exactly the same as in Platonov et al (2019). Patients were seated 70 cm from a liquid crystal display (Dell P2210, resolution 1680 × 1050 pixels, 60 Hz refresh rate) in a familiar environment. The visual stimuli were generated using a personal computer equipped with an open GL graphics card using the Psychophysics Toolbox extensions (Brainard 1997, Pelli 1997) for Matlab (The Math Works, Inc.).

#### Visual stimuli and tasks

The stimuli consisted of 1.167s videos clips showing one of two actors (male or female), standing next to a table and dragging or grasping an object (a blue or red ball) on the table using the right hand. At the start of the video, the hand could be shaped either as a palm or a fist and its position could be either above or on the table. In half of the trials, we increased the size of the video (by 20% of the original) within the aperture. The aperture was created by convolving videos with an elliptic mask causing video clips gradually blur into the black background. Finally, the videos were shown as recorded (actor standing to the right of the table), or inverted around the vertical axis (actor on the left side of the table). These manipulations resulted in 64 (2^6^) videos which were then presented either in the full length (Long trials, Fig1A top) or truncated at time points corresponding to each individual’s 84% action discrimination threshold (ranging from 200 to 250ms, mean 228ms, SD 22ms) or 100ms earlier or later (Short trials, Fig1A bottom). The rest of the movie in the short trials was replaced by a dynamic noise produced by randomly scrambling every pixel in the display on subsequent frames.

All trials started with a baseline period (1s), followed by a variable static phase, created by repeating a first video frame, identical for the two actions, for 283, 458, 583, 733 or 883ms, and then followed by the video displaying either action. If patients could not respond during static and dynamic stimulus presentation, they were given another 2 seconds to reach a decision before the trial ended. As soon as patients pressed a button during any of the 3 trial phases (static frame, video, response epoch), the inter-trial period (1s) started (Fig1A). The trials were organized into 4 blocks of action or gender discrimination tasks (Platonov et al 2019), presented in a fixed order: Action-Gender-Action-Gender. Every block consisted of 2 sub-blocks of 32 trials contained either long or short trials, presented in pseudorandom order. At the beginning of each block, the instruction word ‘action’ or ‘gender’ in Italian was displayed for 5s, instructing patients to perform the two-alternative forced-choice discrimination task by pressing, when ready, a right or left button with the right hand to indicate their decision. In the first 2 blocks the original videos were displayed and in the last 2 the inverted videos. During the trial, a fixation cross was presented near the manipulated object in the center of the screen. During the 1s inter-trial interval, only the fixation cross was visible.

Patients were instructed to fixate the cross in the center of the screen, and the experimenter verified that subjects complied with this instruction. Eye movements were successfully recorded in 18 patients using a noninvasive monitor-mounted infrared video system (SMI iView X 2.8.26) sampling the positions of both eyes at 500Hz. Fixation performance was quantified by the standard deviation of horizontal or vertical eye position per trial, and tabulated as the mean across trials of each condition (Table S3).

#### Preliminary procedures

To familiarize patients with the tasks, they were presented with 2 familiarization blocks of 30 long trials chosen pseudo-randomly such that they contained equal numbers (15) of the 2 actions and the 2 actors. Patients responded by pressing a button and received an auditory feedback indicating either a correct (with low pitch tone) or an incorrect (with a high pitch tone) response. The procedure was repeated until patients made fewer than 2 errors per block. Patients first learned which button corresponded to which actor and then which button corresponded to which action. In addition, every test block was preceded by 10 familiarization trials, reminding patients of the proper button-choices for an upcoming task.

After completing familiarization blocks, patients viewed a block of 30 pseudorandomly chosen trials in which videos were truncated at 3 different time points (150, 250 and 350ms) from video onset with the remaining of the video replaced by dynamic noise. The 84%-threshold was estimated for each patient from these trials and later used in the experiment to create trials in the short condition.

### SEEG Data Recording and Processing

For each implanted patient, recordings started with the selection of an intracranial reference, computed as the average signal of two adjacent leads both exploring white matter. These leads were selected patient-by-patient by clinicians using the following criteria: no response to standard clinical stimulations, including somatosensory (median, tibial, and trigeminal nerves), visual (flash), and acoustical (click) stimulations, no sensory and/or motor behavior evoked by electrical stimulation (Avanzini et al 2016). The stereo-EEG was recorded at 1 kHz with a Neurofax EEG-1100 (Nihon Kohden System).

The recordings from all leads in the gray matter were filtered (band-pass: 0.08–300 Hz; notch: 50 Hz) to avoid aliasing effects and were decomposed into time–frequency plots using complex Morlet’s wavelet decomposition. Power in the gamma (50 to 150Hz) frequency band was extracted in 25ms bins of an interval, extending from 1000ms before the start of the trial to 1000ms after the latest response by the subject, the timing of which differs between subject and task. Following previous intracranial studies (Vidal et al 2010, Platonov et al 2019), gamma power was estimated for 10 adjacent non-overlapping 10-Hz frequency bands, and averaged. The quality of the data was visually inspected using plots of average gamma power in all trials collected for a given condition, to detect the possible presence of IEDs. All trials/channels presenting any IED or other transient electrical artefacts were removed. As shown in Platonov et al (2019) those numbers were relatively small.

The anatomical reconstruction procedures included two basic steps as described in Avanzini et al (2016): 1) identifying the recording leads located in the individual cortical surface using the multimodal reconstructions performed in each patient, and 2) importing these locations into a common template, using the warping of the individual cortical anatomy to the fs-LR average template.

### Statistical analysis

#### Behavior data analysis

Accuracy (% correct) and reaction times (RTs) in the four experimental conditions were computed for each patient. Accuracies in the two long conditions and short gender discrimination were close to perfect performance. In short action discrimination they were, as expected, close to the 84%-threshold. RTs were centered on the mean in all four conditions, except for one patient who had a very small RT (256ms) and low accuracy (63%) for short action discrimination. This subject was removed from the analysis.

#### Stereo-EEG data analysis

The analysis was performed on the average gamma band (50-150 Hz) power sampled with 25ms bins, and z-scored against the 1s baseline period, unless specified differently. Each trial was subdivided into 3 epochs: 1) the 1s baseline epoch, ending with static stimulus onset; 2) the static epoch, defined as the 200ms time window starting 75ms after static onset (283ms was duration of the shortest static phase); 3) the video/dynamic epoch, which started at the end of the static presentation (variable across trials) and lasted until a patient gave a response or the end of the video. This video/dynamic period could be further subdivided into a video and noise period. For each epoch in a trial the mean z-score value was estimated and made available for further analysis such as analysis of variance (ANOVA) with 2×3 or 2×4 designs, using video duration and epochs as factors. When necessary, we corrected for multiple comparison by applying false discovery rate (FDR) correction (Benjamini &Hochberg 1995).

Given the large variability in the time courses of the increases during the dynamic period in short action trials, we developed the following alternative to the ANOVA analysis (using a fixed window) to detect leads with significantly increased activity in short compared to long action trials. We tiled the 1000ms following the video onset with eight 300ms intervals shifted by 100ms, and considered those 300ms intervals ending at most 50ms after the RT in short or long action trials, whichever was shortest. For each of these 300ms intervals we calculated a video enhancement index (vei) as the difference between short and long average z-scored gamma activity divided by the sum of their absolute values. To be considered video-enhanced (VE), leads had to satisfy 3 criteria: 1) at least one 300ms interval had a vei>0.25 (66% increase in short trials), 2) the comparison of average z-scored gamma between short and long trials in the 300ms interval with the highest vei was significant (FDR-corrected, one-tailed t test), and 3) the F value of the interaction between duration of video (3 durations of the truncated videos) and bins (22 bins in 50-600ms interval following video onset) in a 3×22 ANOVA was less than a threshold value chosen below the significance level (F value interaction<1.2, p>0.2) to avoid false negatives. The first two criteria ensured we handled the diversity of timings of the increased activity in the short trials. The last criterion prevented considering off responses to the end of the video as increases in the short trials.

The next step was to distinguish amongst the VE leads those with an increase during the truncated video itself (Early), from those showing an increase later in the dynamic period, during the noise presentation (Late), but allowing for leads to shown an increase over most of the dynamic period (Entire). To this end we used the timing of the composite window made of all (not necessarily contiguous) 300ms intervals with a vei>0.2 (50% increase). Thus we defined *Early-VE* leads as leads in which the composite window started in the first or second 300ms interval and ended in the 3 first intervals. We define *Entire-VE* leads those in which the composite window started in the first two 300ms windows but ended in the fourth 300ms interval or later. Finally *Late-VE* leads were those in which the composite window started in the third interval or later.

The distinction between 3 types of VE leads made a coarse definition of the time course of the significant increase of activity in short compared to long action trials. To investigate the precise timing of the activity increase in short action trials for the individual clusters of any type, we compared short to long trials in each lead in a 100ms sliding window (25ms steps) using one-tailed t test. We calculated for each time interval the proportion of significant leads in each patient, and plotted the mean (across patients) proportion of leads with significant increase as a function of time. We color coded the mean proportion of significant leads in the *significance plots* of single clusters. When grouping clusters on basis of their anatomical location, we plotted the grand mean of proportions of significant leads for the group of clusters as a function of time in *significance curves*. The same procedure was followed to assess the timing of the increase in error compared to correct short action trials.

Given the uniformity of the static-period increases, a fixed window analysis was adequate to define static enhanced leads, unlike the VE leads. Hence we used a post-hoc Tuckey test following the 2×4 ANOVA with duration (long, short) and period (baseline, static, video, noise) as factors.

### Continuous maps

To provide a continuous view of the topographic pattern of specific leads, we built, following Avanzini et al (2016), a circular mask (referred to as a disk) using surface nodes (each lead has exactly 7 nodes) and geodesic distances (i.e., the minimum pathway within the gray matter connecting the source and the target nodes, Van Essen 2012) defined on the fs-LR average template. For each cortical node, we defined the nodes within a 1-cm geodesic distance from the original node and weighted their contribution by a logistic function with unitary amplitude, a steepness of 2 and a midpoint at 7.5 mm. As a result, each node of the cortical mesh was associated with a *disk*, i.e. a collection of surrounding nodes, those within 5 mm of the origin being maximally weighted, and those between 5 and 10 mm gradually reduced in weight, avoiding edge effects.

Using this approach, we computed sampling density and relative responsiveness for each cluster type. Cortical sampling density was calculated as the number of explored leads per disk. Cortical regions with a sampling density lower than 3 leads per disk were masked from subsequent analysis. Relative responsiveness corresponded to the prevalence of a given response pattern amongst responsive leads in a disk, i.e. the number of nodes of a given type of increase in the short action trials (Early-VE, Entire-VE, Late-VE and SE) as a percent of the total number of responsive nodes in a disk. This variable indexes the degree to which a cortical patch responds with a given specific pattern. To ensure reliability, we constrained relative responsiveness maps to nodes whose disk contained at least 2 (Early-VE and SE) or 3 (Entire-VE and Late-VE) leads of that pattern. Each cluster specific map was limited to cortical patches exceeding 250 nodes, and the 3 VE cluster types merged into a single RGB file (Avanzini et al 2018).

These density and relative responsiveness maps were plotted using CARET software (Van Essen 2012) (www.nitrc.org/projects/caret) and directly compared with retinotopic regions defined in Abdollahi et al (2014), with cytoarchitectonic fronto-parietal regions as defined in Avanzini et al (2016), and with the coordinates of the FBA sites (Peelen & Downing 2005).

The multiple demand system was defined as in Fedorenko et al (2013) and its volume made available by J Duncan. We first projected the t-map onto the cortical sheet of fs_LR-average cortical template, and limited the view to regions with a t-score > 1.5, the threshold used in Figure 2 of Fedorenko et al (2013).

## Acknowledgements

The authors are grateful to J Duncan, Cambridge, for making available the MR volume of the multiple demand system.

## List of abbreviations

aFO: anterior Frontal operculum
aIns: anterior Insula
AIP: anterior Intraparietal
aIPS: anterior Intraparietal sulcus
AON: action observation network
aSFS: anterior Superior Frontal sulcus
ATL: anterior temporal lobe
aMCC: anterior Middle Cingulate cortex
BA: Brodmann area
cPFC: caudal Prefrontal cortex
d: dorsal
DIPSA: dorsal intraparietal sulcus anterior
dPFC: dorsal Prefrontal cortex
EBA: extrastriate body area
ECog: electro-corticography
EEG: electro-encephalography
EZ: epileptic zone
FBA: fusiform body area
FCD: focal cortical dysplasia
FDR: false discovery rate
FFA: fusiform face area
fMRI: functional Magnetic Resonance Imaging
FO: Frontal operculum
FO1: cytoarchitectonic area in Orbito-frontal cortex
h: hemisphere
Hz: hertz
IED: interictal epileptic discharge
IFJ: inferior frontal junction
IFS: inferior Frontal sulcus
IPS: Intraparietal sulcus
LO: Lateral occipital
LOTC: lateral occipito-temporal cortex
MD: multiple demand
MEG: magneto-Encephalopgraphy
mIPS: middle intraparietal sulcus
mPreCG: middle Precentral gyrus
MR: magnetic resonance
MT: middle temporal
OFC: orbito-frontal cortex
OMA: observed manipulative action
OP: opercular parietal
OTS: occipito-temporal sulcus
PFC: prefrontal cortex
phAIP: putative human AIP
Phipp: parahippocampus
pInsula: posterior Insula
pIPS: posterior Intraparietal sulcus
PMd: premotor (cortex) dorsal
pMTG: posterior Middle Temporal gyrus
POS: parieto-occipital sulcus
PPC: posterior parietal cortex
Prec: precuneus
pSTS: posterior Superior Temporal sulcus
pTC: posterior Temporal cortex
RT: reaction time
SE: static enhanced
SPL: superior parietal lobule
Stereo-EEG: stereotactic electro-encephalography
TO: Temporal Operculum
V: ventral
VE: video enhanced
vlPFC: ventro-lateral Prefrontal cortex

